# SAM-Dependent Precursor Peptide Engagement by Darobactin Maturase DarE

**DOI:** 10.64898/2026.11.17.767387

**Authors:** Lilly E. Cheek, Sanath K. Kandy, Anthony T. Iavarone, Jarett Wilcoxen, Wen Zhu

## Abstract

Radical *S*-adenosylmethionine (SAM) enzymes that synthesize structurally constrained ribosomally synthesized and post-translationally modified peptides (RiPPs) with therapeutic potential mostly do so by employing a dedicated RiPP recognition element (RRE). With the growing number of RRE-free maturases discovered, how these enzymes engage their substrates remains largely unexplored. In this work, we mapped substrate-enzyme interactions in DarE, an RRE-free radical SAM enzyme that installs the C-C and C-O-C crosslinks of the antibiotic darobactin during the first step of darobactin biosynthesis. Using anaerobic hydrogen-deuterium exchange mass spectrometry (HDX-MS), we simultaneously resolved specific regions within the enzyme and its precursor peptide substrate, DarA, across five complex states. The N-terminal extension and part of the TIM-barrel domain of DarE, along with the leader region of DarA, become protected, revealing the peptide-engagement interface. Additional protection emerged only in the presence of SAM, identifying SAM as a key contributor to precursor-peptide engagement in an RRE-free RiPP maturase. Electron paramagnetic resonance spectroscopy further demonstrated precursor peptide-dependent perturbations of the chemical environment of the iron-sulfur clusters. These findings suggest that SAM-dependent peptide engagement, mediated in part by the N-terminal extension of radical SAM enzyme, provides an alternative mechanism for precursor-peptide recognition in RRE-free RiPP maturases. Our work also demonstrates the utility of anaerobic HDX-MS for characterizing protein-peptide-small molecule interactions in oxygen-sensitive metalloenzymes.

## Introduction

Ribosomally synthesized and post-translationally modified peptides (RiPPs) are natural products with substantial potential for therapeutic discovery.^1–4^ Enzymes that install these unique post-translational modifications are therefore attractive platforms for bioengineering.^5–8^ Among these biosynthetic enzymes, SPASM-domain-containing radical *S*-adenosylmethionine enzymes have emerged as a major class responsible for installing chemically challenging peptide crosslinks that give rise to structurally and functionally diverse RiPPs.^9–16^ Many RiPP-producing radical SAM enzymes contain one or more auxiliary iron-sulfur clusters within the SPASM domain fused to the partial TIM barrel, where the active site [4Fe-4S] cluster is coordinated by the CX_3_CX_2_C motif.^17–20^ Understanding how RiPP-producing radical SAM enzymes recognize and position their peptide substrates is essential for both elucidating their catalytic mechanisms and enabling the enzymatic production of novel RiPPs.^10,21^

A representative example is darobactin, a bicyclic RiPP antibiotic with potent activity against Gram-negative pathogens, including several multidrug-resistant organisms.^22–25^ The first step in darobactin maturation involves C-C and C-O-C cyclization reactions between three amino acid side chains within the precursor peptide DarA, leading to structural rigidification that contributes to its antibiotic effect.^22,24,25^ This reaction is catalyzed by DarE, a SPASM-domain radical SAM enzyme,^23,26^ in the presence of SAM, reductant, and molecular oxygen (Figure 1).^23,26–29^ Biochemical and mechanistic studies on DarE demonstrated key peptide-derived radical intermediates during catalysis and the unusual role of molecular oxygen in this ‘oxygen-sensitive’ enzyme.^26,27,30,31^ The radical chemistry is initiated when the 5’-deoxyadenosyl (5’-dAdo) radical, generated by reductive cleavage of SAM within the partial TIM barrel, abstracts a hydrogen atom from the precursor peptide.^30^ Molecular oxygen is required for the formation of both the C-O-C crosslink between two tryptophan residues and the C-C crosslink between lysine and tryptophan.^26,32^

**Figure 1.**
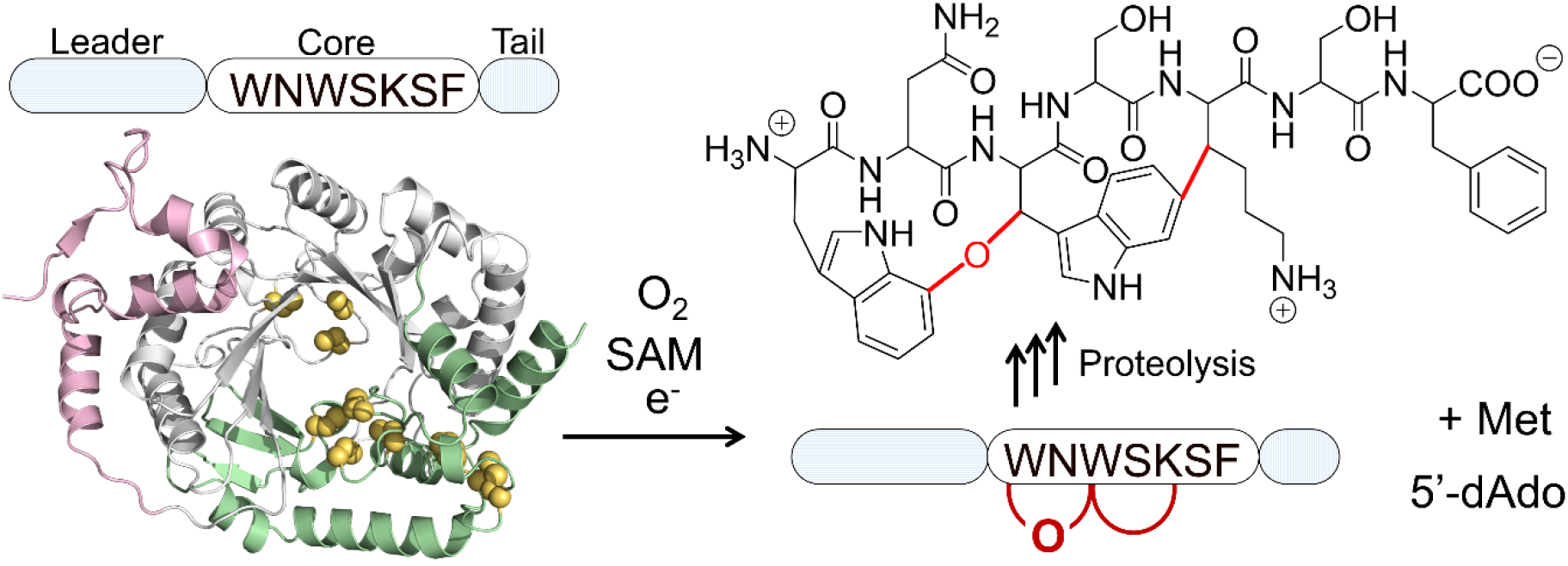
DarE catalyzes the modification of precursor peptide DarA in darobactin biosynthesis. DarA is made of the leader, core, and tail. AlphaFold3-predicted structure of radical SAM enzyme DarE is shown, highlighting the N-terminal extension (pink), radical SAM core domain (gray), and SPASM domain (green). Iron-sulfur clusters are shown as gold spheres. The DarA crosslinking reaction catalyzed by DarE in the presence of oxygen, SAM, and reductant is shown with installed crosslinks highlighted in red.

In addition to the partial TIM barrel and SPASM domain, DarE and other daropeptide maturases contain a conserved N-terminal extension (Figure 1).^31^ DarE, therefore, represents a growing class of RiPP maturases lacking a typical RiPP recognition element (RRE) domain.^33,34^ An RRE domain or protein mediates recognition of the precursor peptide in many well-characterized RiPP radical SAM maturases and has been frequently used to identify novel RiPP gene clusters.^35–39^ RREs typically adopt a winged helix-turn-helix fold and function as peptide chaperones or fusion domains that promote substrate recognition, enhance binding affinity, and facilitate productive enzyme-substrate engagement.^39–41^ Structural studies have shown that the β-sheet of RRE binds to the leader of the precursor peptide, often via backbone hydrogen bonding and/or hydrophobic interactions.^40,42–45^ Although key residues in DarA required for productive cyclization have been identified,^31,46–48^ the regions in DarE responsible for recognizing the leader, core, or tail of DarA segments remain unknown.

Alongside advances in cryogenic electron microscopy and X-ray crystallography for determining structures of enzyme-precursor peptide-SAM complexes,^40,49,50^ hydrogen-deuterium exchange mass spectrometry (HDX-MS) has emerged as a powerful solution-based technique for probing protein-ligand binding and conformational dynamics in recent years, particularly well suited for investigating interactions with intrinsically disordered precursor peptides.^51–54^ In this work, we integrated anaerobic HDX-MS, electron paramagnetic resonance (EPR) spectroscopy, and OpenFold3^55^ modeling to investigate substrate recognition during binary and ternary complex formation that leads to DarA modification. Our study identified recognition sites on DarA and DarE in binary and ternary complexes and highlighted the role of SAM in ternary complex formation. Our solution-based analysis of the protein-peptide interaction provides a foundation for understanding complex formation during the early steps of darobactin biosynthesis and provides a general approach for tackling protein-peptide interactions in otherwise challenging systems.

## Results and Discussion

### An alternative DarE preparation protocol yields a catalytically competent enzyme without chemical reconstitution

As a member of the SPASM-domain-containing radical SAM enzyme subfamily, in addition to the canonical radical SAM [4Fe-4S] cluster (RS cluster) within the partial TIM-barrel, the conserved cysteine-rich iron-sulfur cluster-binding motifs in the SPASM domain are predicted to coordinate two fully ligated auxiliary [4Fe-4S] clusters, AuxI and AuxII (Figure S1a). Therefore, the fully metallocofactor incorporation corresponds to a theoretical total of 12 iron and 12 sulfide ions per DarE monomer.^18,26^ The reported DarE expression and purification protocols required post-purification chemical reconstitution to restore metallocofactor loading.^26,27^ We systematically optimized DarE expression and purification conditions to simplify the protocol (See Materials and Methods in Supplementary Information). Using this alternative preparation protocol, the as-purified DarE sample migrated as a single band on sodium dodecyl sulfate-polyacrylamide gel electrophoresis (SDS-PAGE) and appeared brown in color (Figure S1b). The UV-Vis spectrum of the as-purified DarE exhibited a characteristic absorption feature at 410 nm, indicative of the presence of [4Fe-4S] cluster(s) (Figure S1c). This feature disappeared upon sodium dithionite (DTH) treatment (Figure S1c), consistent with the redox sensitivity of the [4Fe-4S] cluster(s) as previously reported.^17,56,57^ Iron and sulfide quantification revealed 12 ± 1 mol of iron and 14 ± 2 mol of sulfide per monomer, consistent with near-complete occupancy of three [4Fe-4S] clusters. Anaerobic size-exclusion chromatography revealed that DarE exists predominantly as a monomer in solution at concentrations below 10 µM (Figure S1d). At higher protein concentrations, we observed dimeric and oligomeric species, suggesting concentration-dependent oligomerization with an estimated dissociation constant of approximately 50 ± 30 µM for the monomer-dimer transition (Figure S1d). We next evaluated the enzyme activity of the as-purified DarE using high-resolution liquid chromatography-mass spectrometry (LC-MS) assays (Figure S2).^27,54^ Upon incubation of as-purified DarE with DarA, SAM, dioxygen, and the biological reducing system consisting of flavodoxin, flavodoxin NADP^+^ reductase, and NADPH, we observed the time-dependent increasing formation of 5’-dAdo (*m*/*z*=252.10916, [M+H]^+^ ion) and crosslinked DarA peptide products corresponding to doubly crosslinked product (*m/z*=944.19686, [M+7H]^7+^ ion), the C-C crosslink product alone (*m/z=*942.19982, [M+7H]^7+^ ion) and the C-O-C crosslink product alone (*m/z=*944.48481, [M+7H]^7+^ ion), suggesting the presence of the catalytically competent form of the enzyme using this procedure (Figure S2). Fitting the time course of 5’-dAdo production (Figure 2) to a single exponential gave an apparent first-order rate constant of 1.9×10^−3^ min^−1^ and an extrapolated endpoint of 52 µM, approximately one equivalent of 5′-dAdo per DarE. At 18 h only 15% of DarA and 5% of SAM had been consumed, indicating a single slow turnover rather than steady-state catalysis. These results indicated that our as-purified DarE was catalytically competent, with [4Fe-4S] clusters properly incorporated.

**Figure 2.**
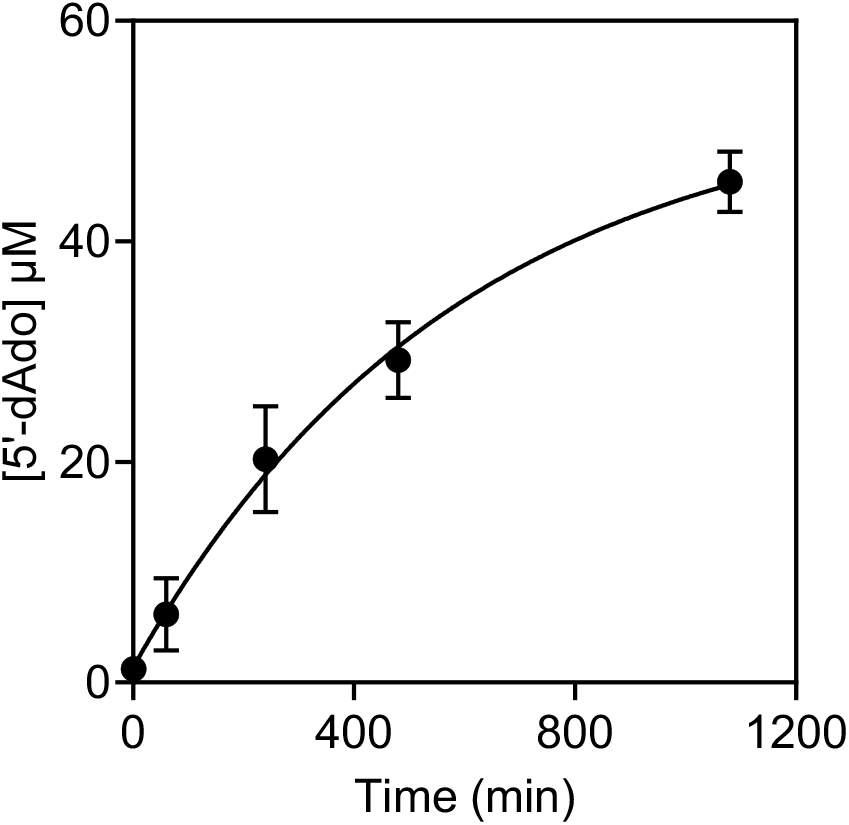
DarE SAM-cleavage activity determined by LC-MS. The product formation curve was fitted to a single-exponential model, giving an apparent first-order rate constant of 1.9×10^−3^ min^−1^ and an extrapolated end point of 52 µM of 5’-dAdo. Points are the mean of two independent biological replicates (Figure S2c).

### HDX-MS identifies regions within DarE and DarA responsible for enzyme-substrate interactions

#### Evaluating experimental setups and establishing comparison methodology

Recent methodological advances have enabled HDX-MS analysis of oxygen-sensitive metalloproteins under anaerobic conditions, providing new opportunities to uncover the protein-peptide interactions in radical SAM enzymes and other oxygen-sensitive enzymes.^54,58–60^ We therefore applied it to explore the protein-peptide interactions in DarE. To achieve high spatial resolution, we optimized the experimental workflow to obtain optimal libraries for both DarE- and DarA-derived peptides under the same conditions. Among the 355 peptides identified by liquid chromatography-tandem mass spectrometry (LC-MS/MS), we selected 24 high-quality, DarE-derived peptides for HDX analysis. This peptide set yields 90.44% sequence coverage and 77.62% amide coverage of DarE (Figure S3a-b). The regions missing from our analysis contain residues 1, 100-102, 144, 180-209, 238-241, and 250-251 (Figure S3a). For DarA, 8 DarA-derived peptides with suitable length and high data quality were selected from the 79 total peptides identified by LC-MS/MS, resulting in 100% sequence coverage and 96.6% amide coverage for DarA (Figure S3c). We subsequently performed time-dependent (t = 0, 45, 600, 1800, 3600, and 10800 s) deuterium-labeling experiments for DarE alone, DarA alone, DarE+SAM (ES) binary complex, DarE+DarA (EA) binary complex, and DarE+SAM+DarA (EAS) ternary complex in duplicates.

We first analyzed the H/D exchange patterns of samples containing DarE alone and DarA alone (Figure S3). The majority of the analyzed DarE-derived peptides clearly exhibited a time-dependent increase in deuterium uptake throughout the six time points, indicating that the labeling time scale used for analysis is sufficient to at least partially capture the exchange kinetics of DarE. In contrast, DarA-derived peptides exhibited a less time-dependent increase in deuterium uptake under the same conditions, suggesting that H/D exchange occurs primarily within 45 seconds. This observation is consistent with the more solvent-exposed, conformationally flexible nature of DarA as the peptide substrate.

To analyze how substrate binding alters H/D exchange in DarE and how interactions with DarE influence the solvent accessibility of DarA within binary and ternary complexes (Figure 3a), we calculated the differences in percentage deuterium uptake (Δ%D) for DarE- and DarA-derived peptides in the absence and presence of their binding partners across all labeling time points (Figure S4 & S5). A positive Δ%D value indicates that ligand binding protects the peptide from H/D exchange. In the notation, the superscript denotes the two states being compared, while the subscript identifies the origin of the peptide. For the DarA-derived peptide, Δ%D_A_ at the first time point (t_1_ = 45 s) was used to identify the H/D exchange protection induced by the binding partner, because exchange occurs within the first 45 seconds, thus it is the most sensitive time point for reporting binding. For the DarE-derived peptide, the summed Δ%D_E_ (ΣΔ%D_E_) values for each peptide were plotted as bar graphs for each pair of comparisons. Comparison of ΣΔ%D_E_ is an empirical method commonly used to highlight consistent differences in deuterium uptake across multiple time points when no substantial differences in exchange kinetics are observed between conditions.^51,61,62^ In addition, an empirical 5% cutoff on each Δ%D was applied to identify DarA- and DarE-derived peptides, with a cumulative ΣΔ%D threshold of 25% across DarE-derived time points, exhibiting meaningful changes under various conditions.^51,61,62^

**Figure 3.**
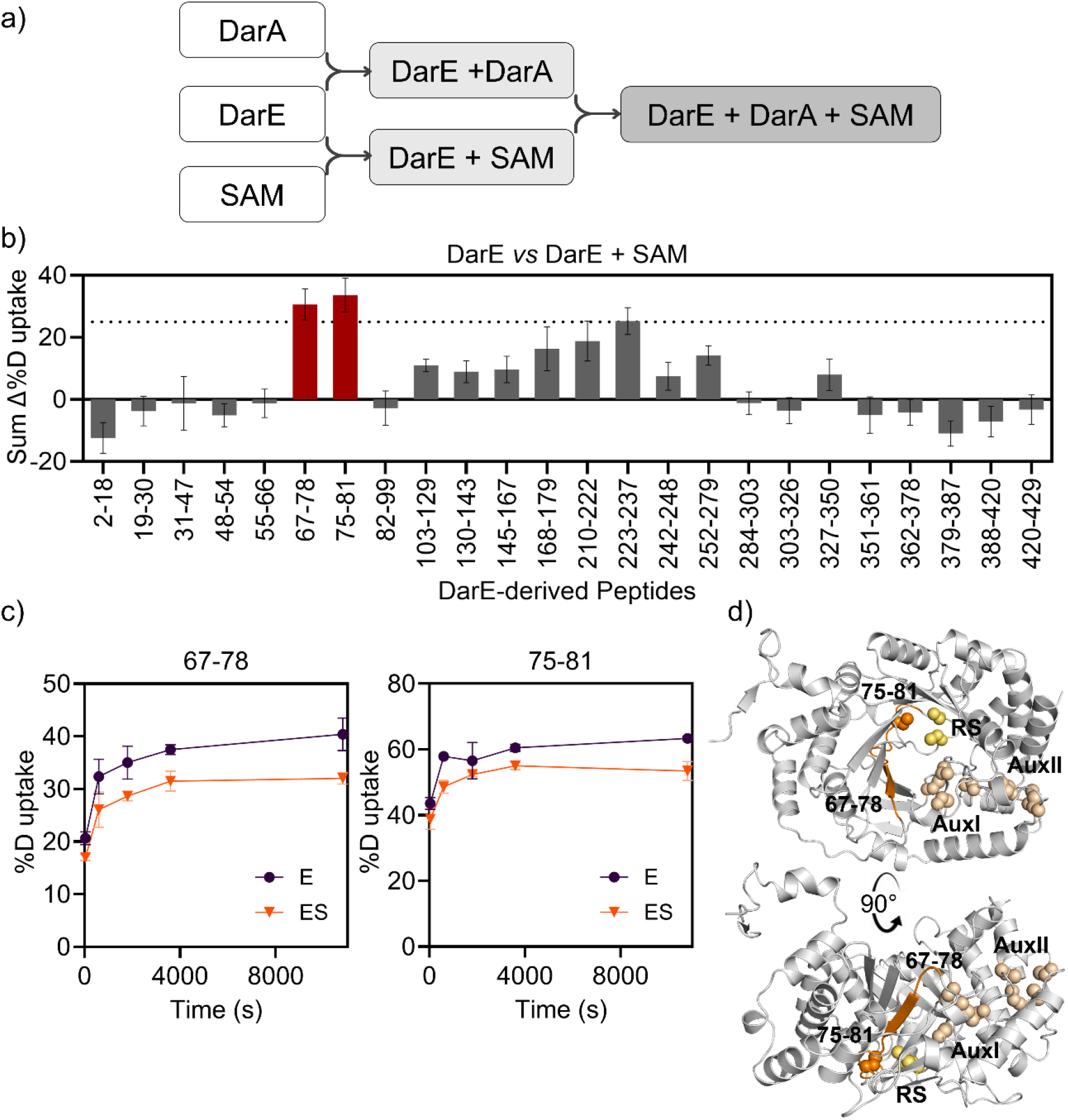
SAM binding protects the radical SAM cluster region of DarE. a) HDX-MS was measured for apo DarE, DarA, the DarE-DarA complex (EA), the DarE-SAM complex (ES), and the ternary DarE-DarA-SAM complex (EAS). b) Sum of differential deuterium uptake (ΣΔ%D_E_) between the DarE-SAM complex (ES) and DarE alone (E) mapped by peptide. Positive Δ%D values indicate protection from hydrogen-deuterium exchange upon SAM binding. Peptides exhibiting notable protection are highlighted in red. Error bars represent the standard deviation of biological duplicate measurements. c) Deuterium uptake plots for p_E_67-78 and p_E_75-81, which display reduced deuterium incorporation upon SAM binding. Deuterium uptake is shown as a function of labeling time for DarE alone (black circles) and the DarE-SAM complex (orange triangles). Data represents the mean ± standard deviation. d) Mapping of SAM-induced protection onto the DarE predicted structural model. Protected peptides p_E_67-78 and p_E_75-81 are highlighted in orange and localize adjacent to the radical SAM (RS) [4Fe-4S] cluster. The cysteines that ligate the RS cluster and the two auxiliary iron-sulfur clusters (AuxI and AuxII) are highlighted in yellow and wheat spheres, respectively.

#### SAM affects the solvent accessibility of the RS site of DarE

We first examined how SAM binding influences DarE H/D exchange behavior in the ES complex. Analysis of the ΣΔ%D_E_^E-ES^ values for each DarE-derived peptide identified two consecutive peptides, p_E_67-78 and p_E_75-81, that exhibited substantial decreases in deuterium uptake, with ΣΔ%D_E_^E-ES^ values of 31 ± 5 and 34 ± 5, respectively, relative to DarE alone (Figure 3b). Time-dependent exchange kinetics showed that protection persisted across all labeling time points (Figure 3c). Since the binding affinity of SAM in radical SAM enzymes is typically in the micromolar range, the experimental conditions used here are expected to maintain DarE predominantly in the SAM-bound state within the ES complex.^63,64^ Therefore, the observed H/D exchange protection most likely arises directly from SAM binding rather than from secondary conformational rearrangements or changes in binding equilibrium during the exchange period.

Peptide p_E_75-81 contains Cys-79, which is part of the highly conserved CX_3_CX_2_C motif responsible for coordinating the RS [4Fe-4S] cluster. Peptide p_E_67-78 partially overlaps with p_E_75-81 and maps to the β_1_-of the partial TIM barrel in the AlphaFold^65,66,66^ predicted structure (Figure 3d & S1a). Therefore, the decreased deuterium uptake upon SAM addition within these peptides indicates that SAM binding reduces solvent accessibility around the RS active site, likely through coordination to the [4Fe-4S] cluster. No other peptides exhibited notable changes in deuterium uptake across the protein under the cutoff criteria applied in this study. Because the SAM-binding mode at the RS [4Fe-4S] cluster has been well established through high-resolution crystallography in similar systems,^40,67–70^ these results provide two important validations of our analysis. First, they demonstrate that the ΣΔ%D_E_ cutoff selected in this study is capable of identifying meaningful ligand-induced protection during H/D exchange without overinterpreting minor changes. Second, they confirm that SAM binding primarily stabilizes the catalytic center as observed in our EPR characterization, and is consistent with observations reported for other SPASM domain-containing RiPP-producing radical SAM enzymes, such as anSME, CteB, PqqE, and SuiB.^40,49,71,72^

#### Identified the interface between DarA and DarE in the EA binary complex

Next, we sought to identify the interface of the enzyme-peptide interaction between DarE and DarA. Comparison of the H/D exchange patterns of the EA complex and DarE alone revealed that two DarE-derived peptides, p_E_168-179 and p_E_210-222, exhibited increased protection in the presence of DarA, with ΣΔ%D_E_^E-EA^ values of 32 ± 8 and 43 ± 7, respectively (Figure 4a & 4b). Both peptides are derived from regions of the partial TIM barrel that correspond to the α_3_β_4_ and α_4_β_5_ structural elements (Figure 4c & S1a). DarA-derived peptides show little or no protection in the EA sample (Figure 4d), indicating that only a minor fraction of DarA is DarE-bound under these conditions and that the flexibility of DarA gave large H/D exchange during dissociation. Our attempt to improve complex occupancy by increasing the DarA concentration resulted in sample aggregation during LC-MS analysis, preventing further enhancement of complex occupancy in solution. Because DarE is expected to be only partially saturated with DarA under the labeling conditions, the observed Δ%D values represent lower bounds on the protection associated with complex formation. The persistent reduction in deuterium uptake for these peptides therefore supports DarA binding at these regions of DarE. In the AlphaFold-predicted structure (Figure 4c), peptides p_E_168-179 and p_E_210-222 are positioned adjacent to one another along the edge of the TIM barrel just above the RS active site, where they likely form a contiguous interaction surface involved in peptide substrate recognition through extensive noncovalent interactions.

**Figure 4.**
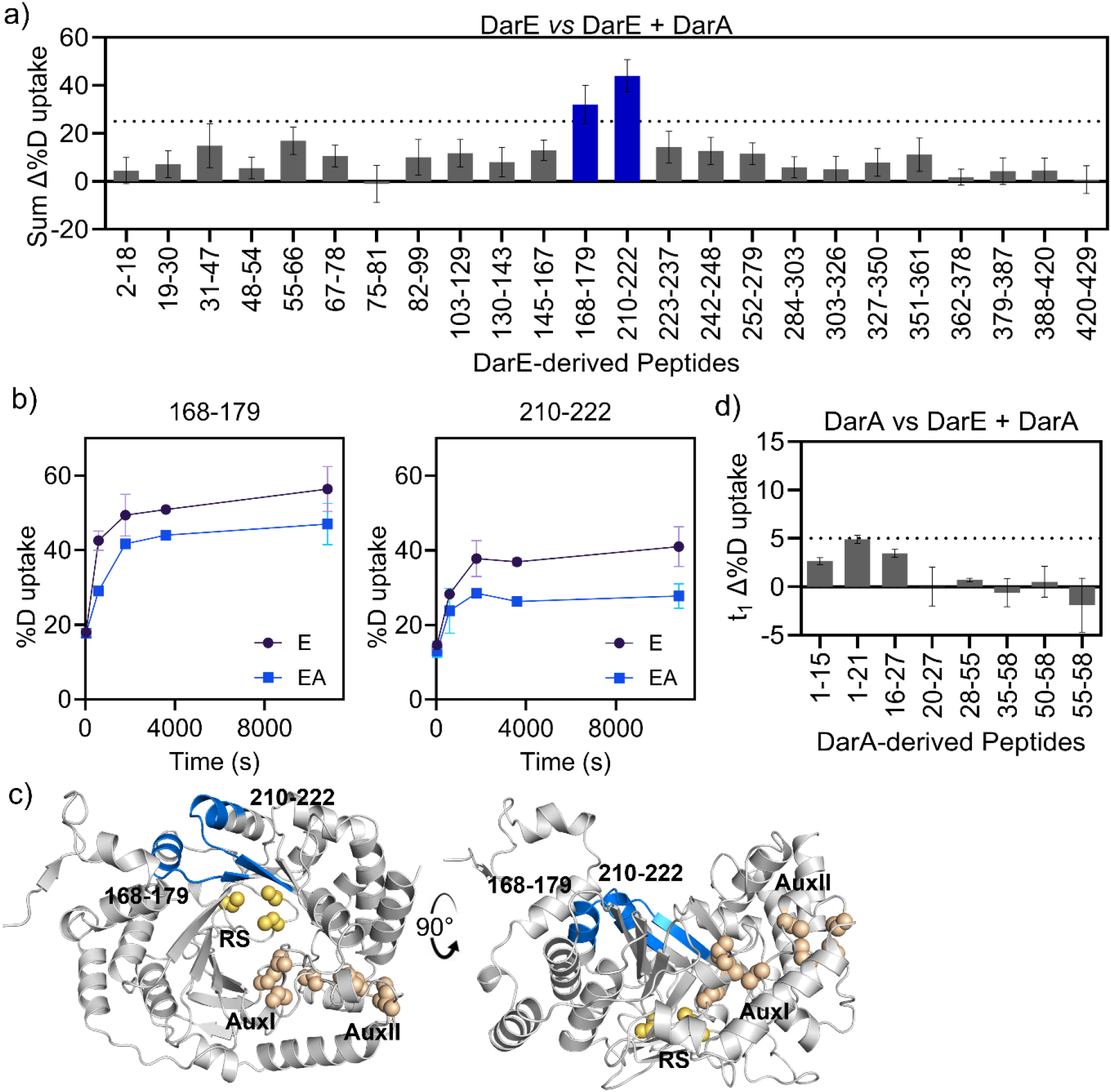
DarA binding protects two regions of the DarE partial TIM barrel. a) Sum of differential deuterium uptake (ΣΔ%D_E_) between the DarE-DarA complex (EA) and DarE alone (E) mapped by peptide. Positive Δ%D values indicate protection from hydrogen-deuterium exchange upon DarA binding. Peptides exhibiting notable protection are highlighted in blue. Error bars represent the standard deviation of biological duplicate measurements. b) Deuterium uptake plots for p_E_168-179 and p_E_210-222, which display reduced deuterium incorporation upon DarA binding. Deuterium uptake is shown as a function of labeling time for DarE alone (black circles) and the DarE-DarA complex (blue squares). Data represents the mean ± standard deviation. c) Mapping of DarA-induced protection onto the DarE predicted structural model. Protected peptides p_E_168-179 and p_E_210-222 are highlighted in blue and localize in the partial TIM barrel. The cysteines that ligate the RS cluster and the two auxiliary iron-sulfur clusters (AuxI and AuxII) are highlighted in yellow and wheat spheres, respectively. d) Differential deuterium uptake analysis of DarA upon DarE addition.

Analysis of the DarA-derived peptides shows no notable Δ%D_A_^A-EA^ upon addition of DarE above the 5% cutoff in the core region (Figure 4d). Only p_A_1-21, derived from the DarA leader, exhibits a borderline Δ%D_A_^A-EA^ of 4.9 ± 0.4, slightly below the cutoff. The lack of notable changes in DarA is consistent with a weak DarA engagement in the absence of SAM.

Combining the observations from these protected regions in DarE and DarA defines a plausible interaction interface between DarA and DarE in the EA binary complex. Peptides p_E_168-179 and p_E_210-222 of DarE potentially interact with the p_A_1-21 of the leader sequence of DarA, anchoring the peptide substrate along the edge of the partial TIM barrel above RS and therefore positioning the C-terminal region of DarA toward the RS active site. No protection was observed for DarE-derived peptides surrounding the RS cluster or for peptides corresponding to the C-terminal core of DarA under these conditions (Figure S5). This observation suggests that, within the EA binary complex, the C-terminal core region of DarA still remains relatively flexible and has not yet fully engaged with the catalytic center.

#### Addition of SAM to the EA complex promotes DarA-DarE interaction

Previous studies of radical SAM RiPP maturases suggest that SAM and precursor peptide can induce additional conformational rearrangements in some systems,^49,54,67,73^ while others show minimal structural changes upon peptide and SAM binding (Table S1).^10,40,74^ To test whether similar behavior exists in DarE, we compared the H/D exchange behavior of DarE-derived peptides in the EAS ternary complex with that in the EA binary complex to determine how SAM binding affects deuterium uptake in DarE and DarA. The same DarE-derived peptides previously assigned to the SAM-binding region, p_E_67-78 and p_E_75-81, were again identified and exhibited even greater protection in the ternary complex, with ΣΔ%D_E_^EA-EAS^ values of 38 ± 4 and 57 ± 6, respectively (Figure 5a). Both of these two values were higher than the corresponding ΣΔ%D_E_^E-ES^ values observed upon SAM binding alone (Figure 3a). The enhanced protection from solvent exchange observed in the ternary complex, therefore, likely arises from direct interactions between DarA and DarE residues within these regions, rather than from a major change in the SAM-binding mode. In addition, peptide p_E_31-47 displayed a ΣΔ%D_E_^EA-EAS^ value of 29 ± 4, modestly exceeding the cutoff threshold applied in this study. Time-dependent exchange plots for these three peptides show that the protected regions consistently exhibit reduced solvent accessibility throughout the labeling period in the EAS complex (Figure 5b). These peptides are derived from the SAM-binding site, along with a short helix in the N-terminal extension unique to DarE, likely reflecting a SAM-dependent influence of DarA binding that was not seen in the EA condition (Figure 5c).

**Figure 5.**
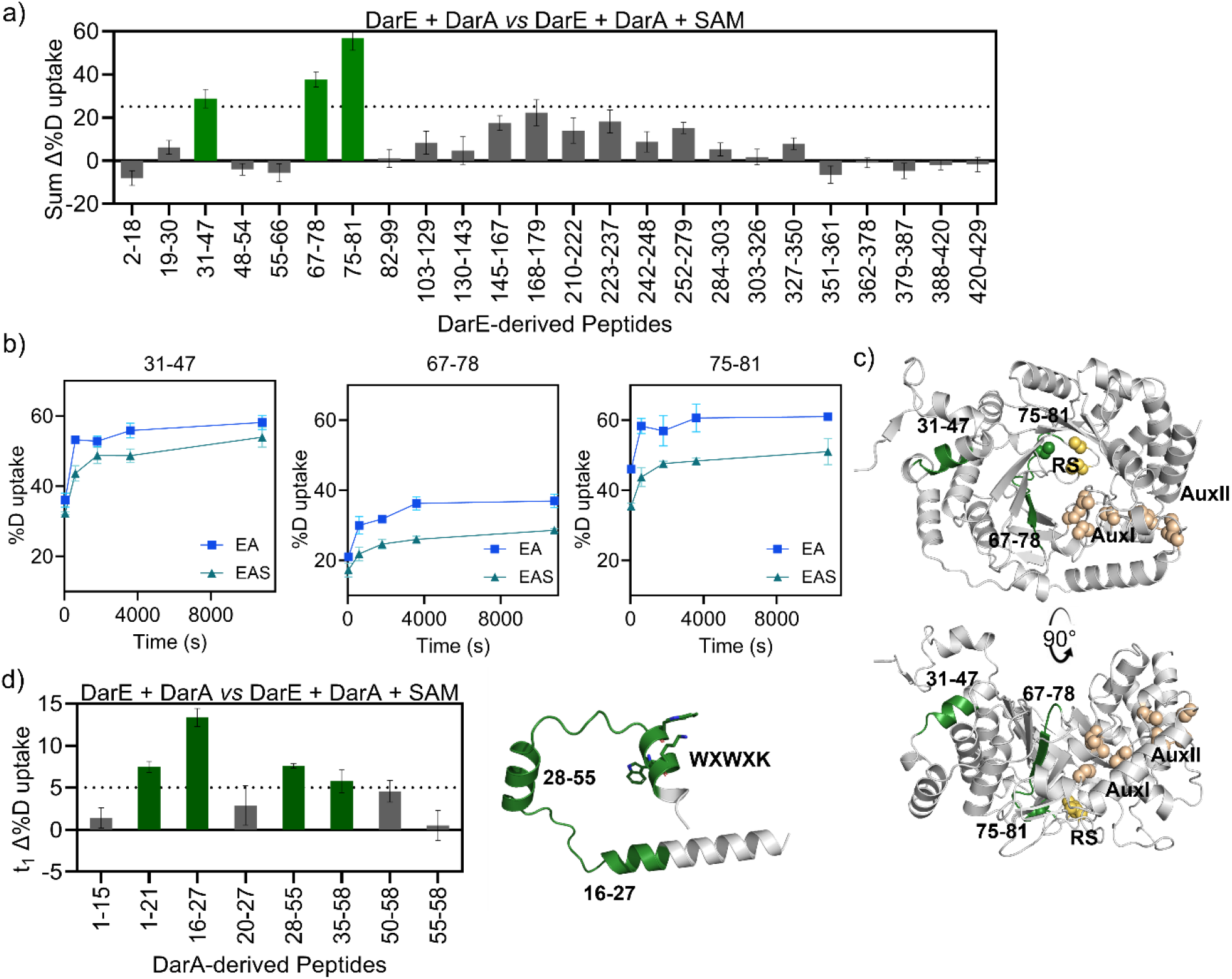
SAM addition to the DarE-DarA complex protects the DarA leader and the DarE N-terminal extension. a) Sum of differential deuterium uptake (ΣΔ%D_E_ between the DarE-DarA-SAM complex (EAS) and DarE-DarA complex (EA) mapped by peptide. Peptides exhibiting notable protection are highlighted in green. Error bars represent the standard deviation of biological duplicate measurements. b) Deuterium uptake plots for p_E_31-47, p_E_67-78 and p_E_75-81, which display reduced deuterium incorporation upon SAM binding. Deuterium uptake is shown as a function of labeling time for EA (blue squares) and EAS (teal triangles). Data represents the mean ± standard deviation. c) Mapping of SAM-induced protection onto the DarE predicted structural model. Protected peptides p_E_31-47, p_E_67-78, and p_E_75-81 are highlighted in green and localized in the N-terminal extension and the SAM binding site within the partial TIM barrel. The cysteines that ligate the RS cluster and the two auxiliary iron-sulfur clusters, AuxI and AuxII, are highlighted in yellow and wheat spheres, respectively. d) Left: Differential deuterium uptake analysis of DarA-derived peptide upon SAM addition to EA complex. Peptides p_A_1-21, p_A_16-27, p_A_28-55, and p_A_35-58 exhibit notable protection upon DarE binding and are highlighted in green. Right: Peptide p_A_16-27, located in the N-terminal leader of DarA, residues 28-50, as shown in the AlphaFold model. The core sequence WXWXK is highlighted in sticks.

Among the peptides derived from DarA, four peptides, p_A_1-21, p_A_16-27, p_A_28-55, and p_A_35-58, have increased protection in the EAS complex compared to the EA complex, with Δ%D_a_^EA-EAS^ values of 7.5 ± 0.7, 13 ± 1, 7.6 ± 0.3, and 6 ± 1, respectively (Figure 5d). These four peptides span most of the DarA sequence, indicating protection across the peptide at early labeling times in the EAS complex. Since p_A_1-21 was the only one that exhibited a modest Δ%D_A_^A-EA^, the further increase throughout the entire DarA observed in the Δ%D_A_^EA-EAS^ suggests that the additional tightening of this region was a result of SAM binding stabilizing the engagement between DarA and DarE. Removing the overlapping peptides that show no notable Δ%D_A_^EA-EAS^ or Δ%D_A_^A-EA^ further refines the region that primarily contributes to the increase in Δ%D_A_^EA-EAS^ to residues 16-55. Within this region, Δ%D_A_^EA-EAS^ of p_A_16-27 experiences the highest increase when compared to that of the Δ%D_A_^A-EA^ (Figure 4d).

Intriguingly, analyses of the DarA sequence alignment showed that p_A_16-27 is part of the only region in the leader sequence that is highly conserved among different darobactin-like precursor peptides (Figure S6). The importance of the leader sequence has also been validated by truncations in which deletion of the first 20 residues lost the ability to be modified by DarE, and additional alanine mutations on highly conserved residues located within p_A_16-27, such as S17A, L18A, and F22A, had a notable reduction in crosslink product formation.^46^ On the other hand, the identified DarE-derived peptide p_E_31-47 also occupied a conserved region in the DarE N-terminal extension (Figure S7). The AlphaFold-predicted model suggests this region contains several helices that likely interact with α1, α2, and α3 of the partial TIM barrel (Figure S1a). One plausible explanation for the simultaneously increased protection in the leader of DarA and N-terminal extension of DarE is that the formation of the ternary complex does not substantially alter the binding mode of either the SAM or the peptide substrate but instead increases DarA occupancy or residence time via SAM-DarA interactions, thereby increasing ligand residence time within the same binding pocket and leading to greater protection from exchange. Alternatively, formation of the ternary complex may induce changes in the binding mode of one or more ligands, including DarA and/or SAM, resulting in additional intermolecular interactions and further amide protection. Both scenarios are consistent with the observed HDX-MS data and cannot be distinguished solely by the current measurements. Future experimental structural validation is therefore necessary to distinguish these two plausible mechanisms.

#### Interaction between DarA and DarE is modulated by SAM

When we analyzed the ΣΔ%D_E_^ES-EAS^ values obtained from the comparison of the EAS and ES complexes to examine the influence of DarA binding on DarE in the presence of SAM, the two peptides previously assigned as the DarA-binding interface, p_E_168-179 and p_E_210-222, again exhibited reduced deuterium uptake, with ΣΔ%D_E_^ES-EAS^ values of 42 ± 6 and 27 ± 4, respectively (Figure 6a). Relative to the corresponding ΣΔ%D_E_^E-EA^ values of 32 ± 8 and 43 ± 7, these modest differences may reflect localized changes in interaction mode between DarA and DarE at the DarA binding site or altered solvent accessibility due to the presence of the SAM. The persistent protection observed for these two peptides across multiple comparisons further supports their assignment as part of the DarA-binding interface in DarE.

**Figure 6.**
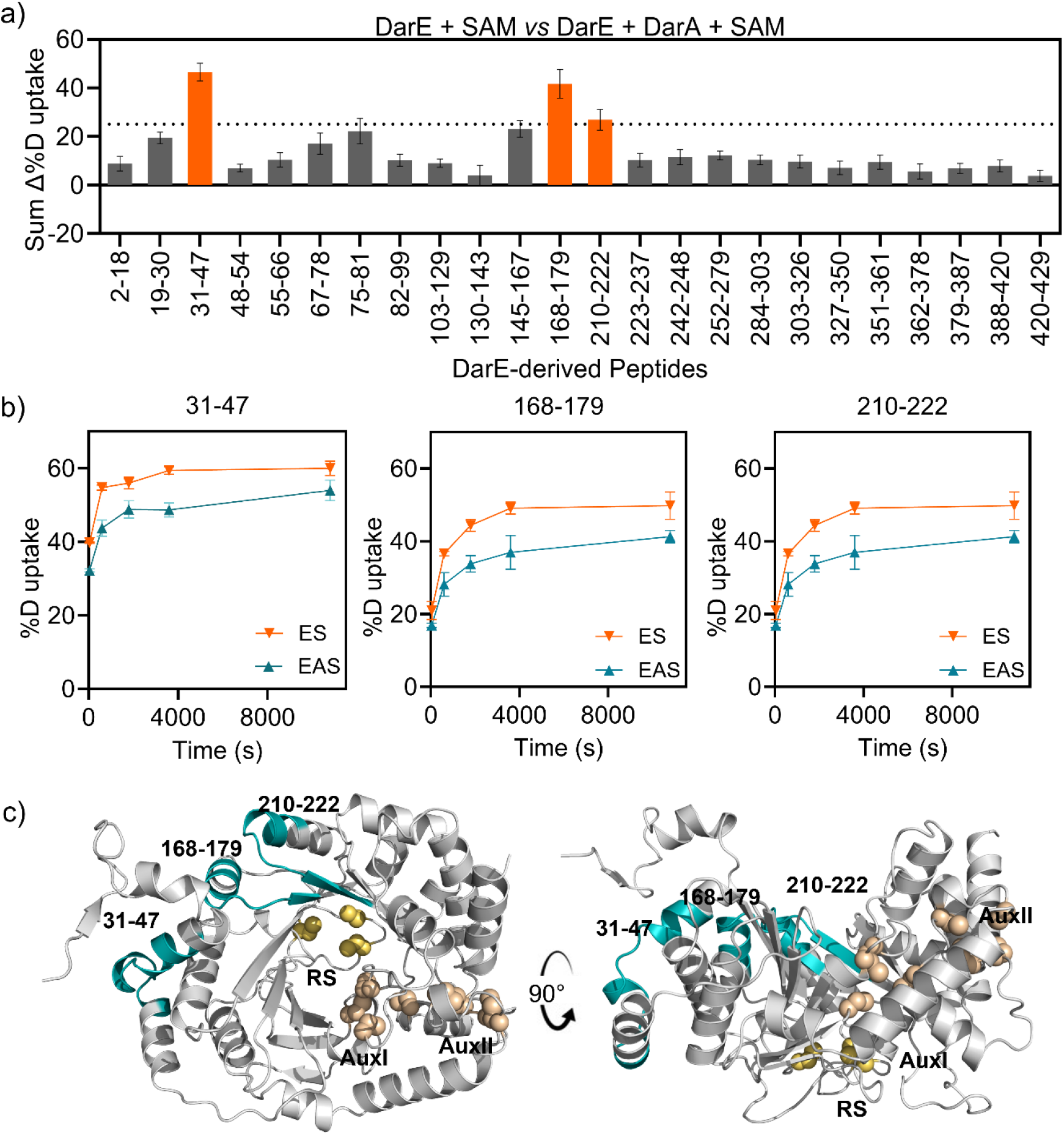
DarA addition to the DarE-SAM complex protects the N-terminal extension and partial TIM barrel. a) Sum of differential deuterium uptake (ΣΔ%D_E_) between the DarE-DarA-SAM complex (EAS) and DarE-SAM complex (ES) mapped by peptide. Peptides exhibiting notable protection are highlighted in orange. Error bars represent the standard deviation of biological duplicate measurements. b) Deuterium uptake plots for p_E_31-47, p_E_168-179 and p_E_210-222, which display reduced deuterium incorporation upon DarA binding. Deuterium uptake is shown as a function of labeling time for ES (orange triangle) and EAS (teal triangles). Data represents the mean ± standard deviation. c) Mapping of DarA-induced protection onto the DarE predicted structural model. Protected peptides p_E_31-47, p_E_168-179 and p_E_210-222 are highlighted in teal and localized in the N-terminal extension and the partial TIM barrel. The cysteines that ligate the RS cluster and the two auxiliary iron-sulfur clusters (AuxI and AuxII) are highlighted in yellow and wheat spheres, respectively.

Once again, we observed that peptide p_E_31-47, located within the N-terminal extension of DarE, displayed a ΣΔ%D_E_^ES-EAS^ value of 42 ± 4, representing the largest protection observed in this comparison set (Figure 6a & 6b). This peptide was also identified in the ΣΔ%D_E_^ES-EAS^ analysis (Figure 5a & 5b) and was previously assigned as a region undergoing DarA-induced local conformational stabilization in the ternary complex. These observations suggest that formation of the ternary complex further stabilizes regions outside the immediate DarA-binding interface at the N-terminal extension of DarE (Figure 6c).

#### Comparing EAS with DarE or DarA alone identifies additional peptides with subtle changes

When directly comparing the H/D exchange patterns of the EAS complex and DarE alone (Figure 7a), we further identified three additional DarE-derived peptides, p_E_145-167, p_E_223-237, and p_E_252-279, that exhibited notable protection with ΣΔ%D_E_^E-EAS^ values of 31 ± 4, 33 ± 4, and 25 ± 3, respectively. All identified peptides displayed similar exchange kinetics under both conditions (Figure 7b), indicating that the observed differences primarily reflect persistent changes in solvent accessibility. Mapping these protected regions onto the structural model provides a clear visualization of the overall H/D exchange protection induced by ternary complex formation, with the majority of protected regions localized to the partial TIM barrel and the N-terminal extension of DarE (Figure 7c). In addition to the previously mentioned peptides, p_E_252-279 emerged in this comparison. This peptide is primarily comprised of a long loop in the partial TIM barrel (Figure 7c). Interestingly, similar loop regions have been observed in other RiPP-modifying radical SAM enzymes, including SuiB,^49^ in which substrate binding is associated with localized conformational rearrangements around the catalytic cavity. Given that HDX-MS data primarily support localized changes in solvent accessibility and do not, on their own, establish the magnitude or directionality of these conformational rearrangements, our working model is that binding of SAM and DarA binding stabilize the loop region near the RS site since such changes were not observed in the binary comparisons.

**Figure 7.**
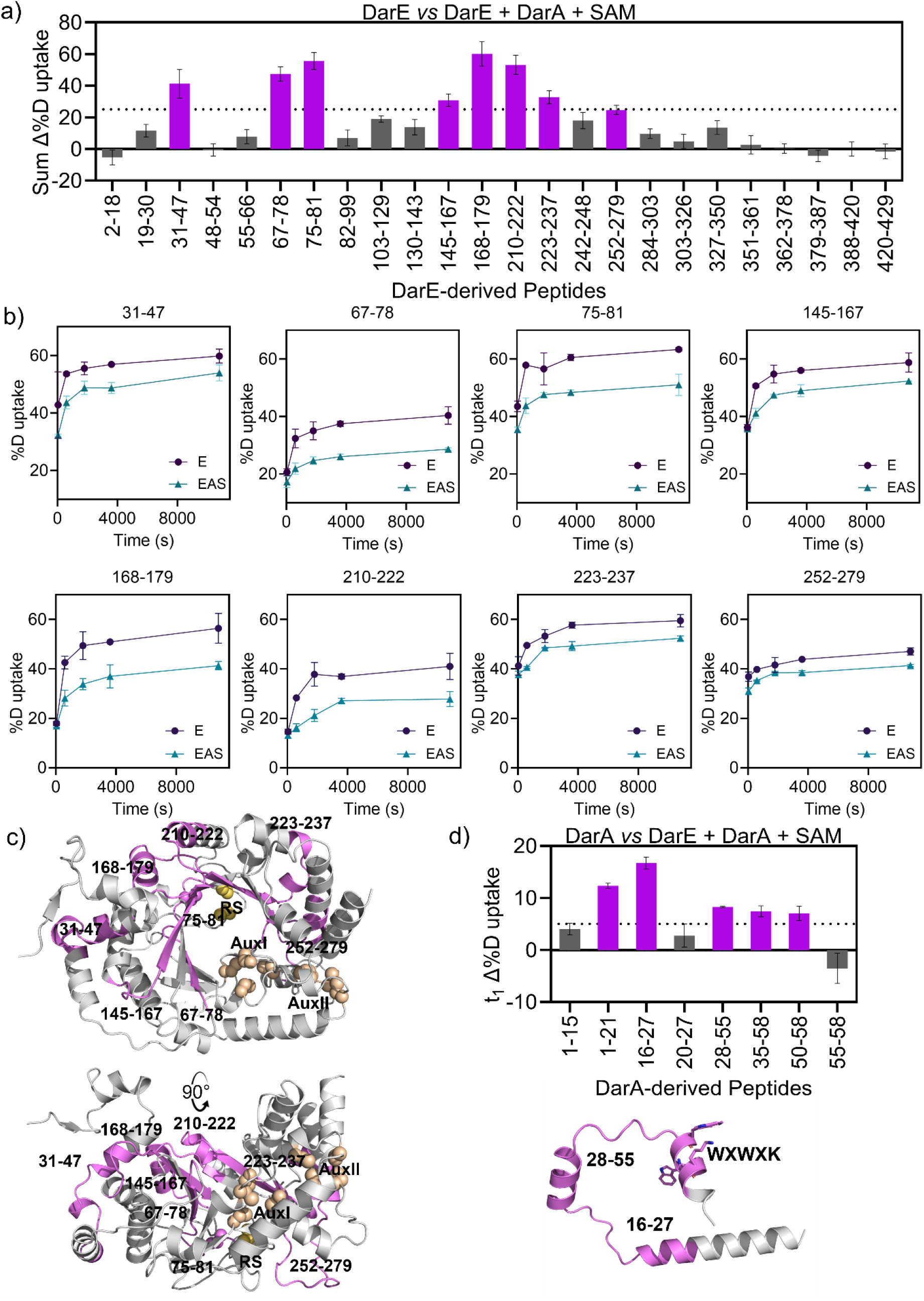
The ternary complex combines the protection observed in both binary complexes. a) Sum of differential deuterium uptake (ΣΔ%D_E_) between the DarE-DarA-SAM complex (EAS) and DarE (E) mapped by peptide. Peptides exhibiting notable protection are highlighted in violet. Error bars represent the standard deviation of biological duplicate measurements. b) Deuterium uptake plots for p_E_31-47, p_E_67-78, and p_E_75-81, p_E_145-167, p_E_168-179, p_E_210-222, p_E_223-237, and p_E_252-279, which display reduced deuterium incorporation upon SAM binding. Deuterium uptake is shown as a function of labeling time for E (blue squares) and EAS (teal triangles). Data represents the mean ± standard deviation. c) Mapping of DarA-induced protection onto the DarE predicted structural model. Protected peptides p_E_31-47, p_E_67-78, and p_E_75-81 are highlighted in green and localized in the N-terminal extension and the SAM binding site within the partial TIM barrel. The cysteines that ligate the RS cluster and the two auxiliary iron-sulfur clusters (AuxI and AuxII) are highlighted in yellow and wheat spheres, respectively. d) Top: Differential deuterium uptake analysis of DarA-derived peptide upon SAM and DarA addition to DarE. Peptide p_A_1-21, p_A_16-27, p_A_28-55, p_A_35-58, and p_A_50-58 exhibit notable protection and are highlighted in violet. Bottom: Peptide p_a_16-27, located at the N-terminal leader sequence region of DarA and residues 28-50, as shown in the AlphaFold model. The core sequence WXWXK is highlighted in sticks.

For the DarA-derived peptides, one additional overlapping peptide, p_A_50-58 (Δ%D_A_^A-EAS^ = 7 ± 1), was identified besides the four peptides that were previously identified, p_A_1-21, p_A_16-27, p_A_28-55, and p_A_35-58 in the EA-EAS comparison (Figure 7d). Peptide p_A_35-58 covers the majority of the conserved sequence (Figure S6), including the ITA motif near the core region, which has been proposed to help position the substrate.^46^ Residues 50-58 contain the partial core peptide sequence WXWXKSF (residue 49-55) and the tail (residues 56-58). Since the Δ%D_A_^A-EAS^ of the tail p_A_55-58 is below our cutoff, the tail contribution to the H/D exchange difference observed should be minor. This is also consistent with the sequence alignment, in which the tail sequence varies across organisms (Figure S6).

### Engagement of SAM and DarA in DarE affects the iron-sulfur clusters as revealed by EPR

To probe the electronic environment of the iron-sulfur clusters in DarE and assess the influence of substrate binding, continuous-wave (CW) EPR spectra were collected for DTH-reduced DarE in the presence of SAM and/or DarA. Reduced DarE alone is expected to contain overlapping signals from up to three [4Fe-4S]^1+^ clusters if their reduction potentials fall within the range that can be reduced by DTH.^75,76^ Consistent with this expectation, the EPR spectrum of DarE alone revealed at least two distinct paramagnetic species, corresponding to two of the three Fe-S clusters present in the enzyme with *g*-values of [2.045, 1.920, 1.850] and [1.9751, 1.9841, 1.9147] for auxiliary cluster A and cluster B, respectively (Figure 8a). We note an unusually low *g_1_* for a typical [4Fe4S] cluster in cluster B, this may be a result of a broadened and difficult-to-detect *g_1_* feature in the EPR, or a result of the unusual Cys-Cys motif, C413-C414, of the AuxII Fe sites. Consequently, cluster A has a higher level of apparent reduction with a larger contribution of the EPR signal across all samples examined, with the largest difference in intensity in DarE alone (Table 1).

**Figure 8.**
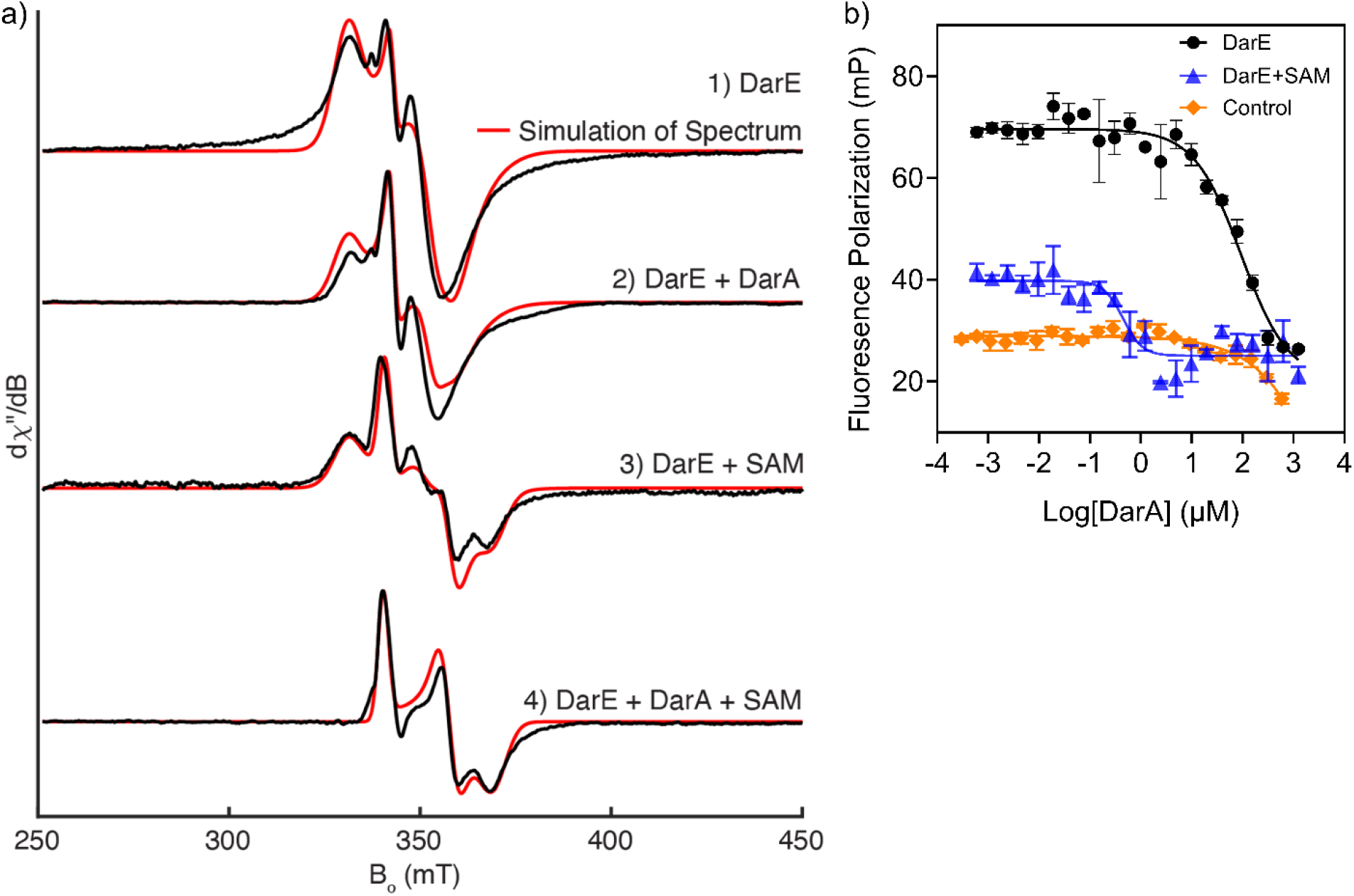
Characterization of DarE-DarA interaction. a) Continuous-wave X-band EPR spectrum of anaerobically purified DarE and various substrate-bound forms. The spectrum is characteristic of reduced [4Fe–4S]^+^ clusters and was collected under the indicated experimental conditions. Simulated spectra are overlaid where appropriate. b) Fluorescence competitive assay of DarE with FITC-labeled DarA in the presence of unlabeled DarA. Increasing concentrations of unlabeled DarA were titrated into a fixed concentration of FITC-labeled DarA-DarE complex in the absence of SAM (black circles) or in the presence of SAM (blue triangles). The control (orange diamonds) omits DarE from the mixture. Points are mean ± SD of triplicate wells.

**Table 1.** Simulation Parameters used for the EPR spectrum of DarE and its complexes.

| *g*-tensor | RS Cluster<br>[1.98836, 1.89177, 1.833] | Cluster A<br>[2.045, 1.920, 1.850] | Cluster B<br>[1.9751, 1.9841, 1.9147] |
|---|---|---|---|
| DarE | 0% | 95% | 5% |
| +SAM | 38% | 58% | 4% |
| +DarA | 0% | 88% | 12% |
| +DarA+SAM | 100% | 0% | 0% |

Addition of SAM to reduced DarE resulted in an overall weakening and broadening of the EPR features observed in the sample with DarE alone and was accompanied by the appearance of a new species between 340 and 370 mT (Figure 8a). Differential analysis of the spectra of DarE alone and DarE in the presence of SAM identified a single dominant species with *g*-values of [1.98836, 1.89177, 1.833], which we assign to the radical SAM (RS) cluster (Figure 8a & S8). Observation of the RS cluster only upon SAM binding has also been reported for other radical SAM enzymes, where coordination of SAM to the unique iron of the [4Fe-4S] cluster alters the electronic structure and stabilizes the EPR-active reduced state.^64,77^ The RS [4Fe-4S]^1+^ cluster typically exhibits axial or slightly rhombic EPR signals centered near *g* = 2.04, arising from the *S* = 1/2 ground state of the reduced cluster (Table S2).^76,78–80^

Incubating DarA with reduced DarE produced a substantially more intense EPR signal compared to DarE alone, with an increase in the contribution of cluster B to the spectrum (Figure 8a). Like DarE alone, the signals corresponding to the two auxiliary Fe-S clusters are the only species observed, with the RS cluster remaining unreduced in the absence of SAM or significantly broadened, impairing detection (Table 1). These observations indicate that peptide binding significantly perturbs the electronic environment of the auxiliary Fe-S cluster network within DarE. The increased contribution of cluster B to the overall spectrum relative to cluster A may reflect changes in spin coupling, reduction potential, or conformational dynamics.

The EPR spectrum of the ternary complex containing DarE, SAM, and DarA is distinct from that of either binary complex, with loss of the EPR spectrum associated with the auxiliary clusters and enhancement of the RS cluster EPR species (Table 1 & Figure 8a). These ligand-dependent changes indicate that the electronic environment of the clusters differs in the ternary complex from either binary complex, consistent with reorganization near the radical SAM cluster upon substrate and cofactor binding. Furthermore, peptide engagement in radical SAM enzymes is expected to be coupled with RS active-site reorganization, and these ligand-dependent spectral changes are consistent with DarA-induced remodeling of the catalytic pocket surrounding the RS [4Fe-4S] cluster.^49,54^ Similar ligand-dependent perturbations of Fe-S cluster EPR features have been observed in other radical SAM enzymes (Table S2) and are frequently associated with activation of the organometallic intermediate-forming state preceding reductive SAM cleavage.^81–83^

Recent EPR studies revealed that hydrogen abstraction from the Cβ position of Trp51 in the core sequence of DarA initiates the radical-based catalytic cascade responsible for double crosslink formation, supporting a proposed chemical mechanism.^30^ The functional roles of the auxiliary Fe-S clusters in DarE remain unclear. We therefore attempted to generate single- and double-cluster knockout variants by systematically mutating the cluster-ligating cysteine residues to alanine. All variants, however, exhibited severe stability defects during expression and purification, preventing further spectroscopic characterization and unambiguous assignment of the auxiliary cluster signals (clusters A and B) to the expected AuxI or AuxII clusters identified by sequence and structural similarities to other SPASM domain enzymes. Although the experimental spectra can be reasonably reproduced using a three-cluster model, only the RS cluster could be confidently assigned in this study based on its SAM-dependent spectral features. Identification of the RS cluster enabled simulation of the composite EPR spectra using three contributing iron-sulfur cluster species corresponding to the RS, AuxI, and AuxII clusters (Figure 8a & Figure S8). In these simulations, only the relative weights of the individual cluster contributions were varied across samples, with the simulation parameters for each cluster held constant (Table 1). As described above, the variable intensities of the individual Fe-S clusters under the various substrate conditions, under which the spectra were collected, likely reflect changes in the reduction potential and/or conformational dynamics near the clusters.

### SAM alters DarA-DarE interaction in a fluorescence polarization competition assay

We performed a fluorescence polarization (FP) competition assay to further probe the role of SAM in the interactions between DarA and DarE. An N-terminal fluorescein-labeled DarA peptide was used as the probe, and unlabeled DarA was titrated anaerobically into DarE in the presence and absence of SAM. Titration curves acquired in the presence of SAM were displaced toward lower DarA concentrations than those acquired without SAM (Figure 8b), indicating that SAM promotes DarA engagement. In the absence of DarE, the DarA titration decreased the FP signal at high concentrations of DarA, which prevented us from assigning a quantitative IC_50_ for DarA-DarE interactions in the presence and absence of SAM. For radical SAM RiPP maturases that contain a fused RRE domain, peptide-enzyme interactions typically occur with nanomolar to low-micromolar affinities.^10,39,84^ The RRE generally provides an extended binding surface that recognizes the leader peptide through hydrophobic and hydrogen-bonding interactions, thereby increasing precursor peptide affinity.^13,35,41^ For example, the radical SAM enzyme PqqE exhibits no detectable interaction with its precursor peptide PqqA in the absence of the standalone RRE protein, PqqD.^84^ Formation of the PqqD-PqqA complex enhances binding to PqqE, resulting in an apparent affinity of approximately 0.2 μM.^84^ SAM may therefore promote substrate engagement in DarE, partially compensating for the absence of a dedicated peptide-recognition domain.

### Integrating experimental observation and OpenFold3 to define the DarA-DarE interaction interface

To integrate the HDX-MS-identified regions involved in substrate engagement with structural and sequence information, we compared our HDX-MS results with sequence conservation and an OpenFold3^85,86^ model of the EAS complex. Among the five top-ranked models, the overall architecture of the complex was similar, with the primary structural variation occurring in the conformation of the C-terminal tail of DarA (Table S3 & Figure S9). In the absence of additional structural restraints, we selected the top-ranked model for subsequent analysis and illustration (Figure S9 & Figure S7).

In the predicted EAS complex, SAM is positioned adjacent to the radical SAM [4Fe-4S] cluster in a configuration consistent with the canonical SAM-binding mode of characterized radical SAM enzymes and with our experimental observations (Figure 9). The DarA core resides near the conserved radical SAM CX_3_CX_2_C motif, coinciding with enhanced protection of peptides p_E_67-78 and p_E_75-81 was observed in the ternary complex relative to the ES complex. Several DarE regions that exhibited comparatively subtle but reproducible protection of H/D exchange in the presence of both substrates, including peptides p_E_145-167, p_E_210-222, p_E_223-237, and p_E_252-279, are also predicted to interact at or near the DarA core in the OpenFold3 model (Figure 9 & Figure S9). The predicted architecture agrees with the HDX-MS protection pattern and places the DarA core in proximity to regions of DarE that become specifically stabilized in the catalytically relevant ternary complex.

**Figure 9.**
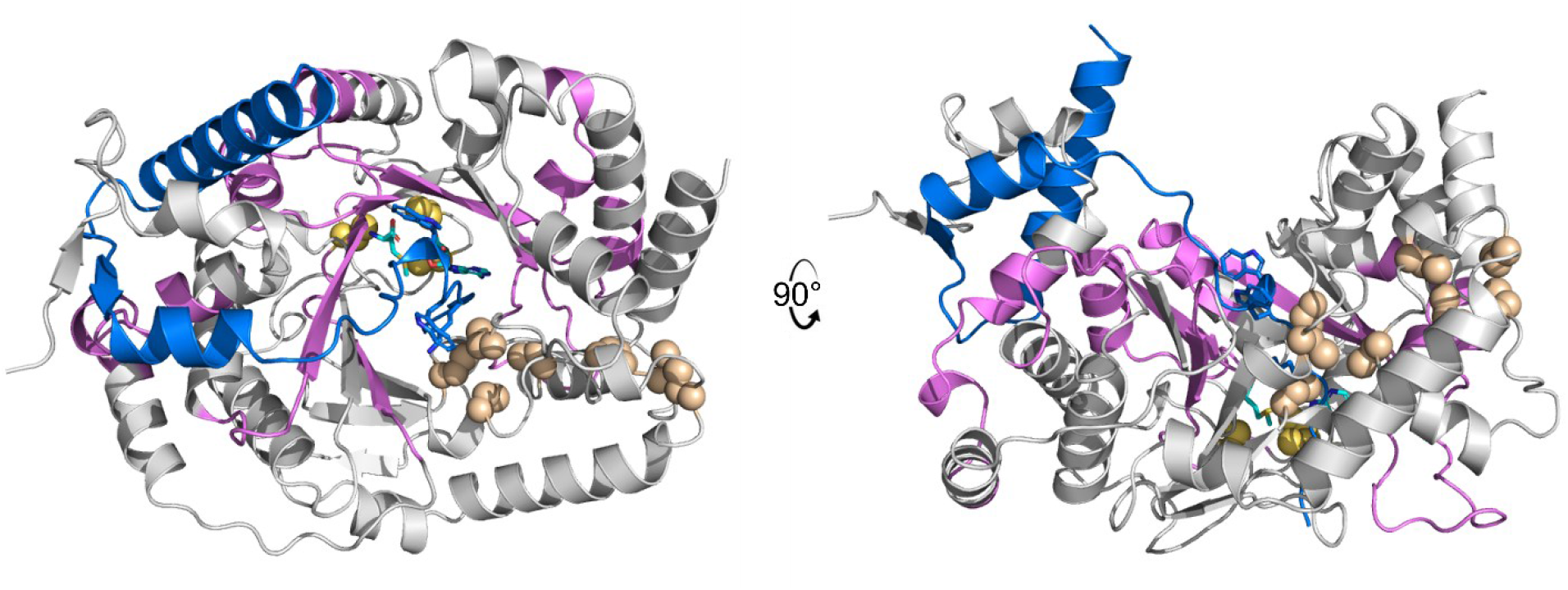
HDX-MS-identified interactions mapped onto the OpenFold3-predicted EAS complex. DarA is shown in blue. The conserved DarA core motif, WXWXK, is shown as sticks. DarE regions identified by HDX-MS as contributing to the formation of the EAS complex are highlighted in pink; the remainder of DarE is shown in gray. SAM is shown as cyan sticks. Cysteines coordinating the three [4Fe-4S] clusters are shown as spheres, with the radical SAM (RS), auxiliary I (AuxI), and auxiliary II (AuxII) clusters colored yellow, wheat, and wheat, respectively.

The predicted interface also provides a structural framework for understanding recognition of the DarA leader peptide. In the OpenFold3 model, the leader contacts the N-terminal extension of DarE, p_E_31-47, as well as α3β3 and α4β4 of DarE, including p_E_145-167 and p_E_168-179. (Figure S9 & S1A) Previous mutagenesis studies identified conserved DarA residues S17, L18, and F22 as important for productive interaction with DarE.^46^ The same study also proposed that conserved residues S17, L18, and F22 in the DarA leader facilitate the DarE binding.^46^ Our analysis further supports that hydrophobic regions within DarE, particularly p_E_31-47 and p_E_168-179, contribute to recognition of this portion of the leader peptide.

Sequence conservation provides additional evidence supporting the key regions present in both DarA and DarE interface (Figure S6 & S10). The protected DarE regions contain conserved hydrophobic motifs, including an IIGXI sequence within p_E_31-47 and a WLVFLKGI motif within p_E_168-179, which are positioned in the OpenFold3 model near hydrophobic residues of DarA, including L15, L18, F22, and I29 (Figure 9). The DarE N-terminal extension is similarly enriched in conserved hydrophobic residues (Figure S10), suggesting that hydrophobic interactions may provide an important energetic contribution to leader recognition. The convergence of sequence conservation, HDX-MS protection, and the predicted structural interface therefore supports a model in which the N-terminal extension of DarE mimics the role of the RRE domain in engaging DarA, likely through primarily hydrophobic interactions, and DarA binding further stabilizes the radical SAM active-site region for catalysis. Meanwhile, the auxiliary iron-sulfur clusters and conserved regions in the SPASM domain (Figure S11) may influence catalysis primarily through electronic effects, as seen in the EPR analysis, rather than extensive protein-substrate contacts within the auxiliary domain environment since no peptides within the SPASM domain showed protection above the cutoff in this study.

## Conclusion

Using a combination of solution-based approaches, we characterize the interaction between DarE and its precursor peptide, DarA, during the early step of darobactin biosynthesis. Our time-dependent anaerobic HDX-MS identified regions of DarE and DarA that exhibit altered solvent accessibility upon formation of binary and ternary complexes containing SAM and the peptide substrate. Additionally, EPR suggests that the redox state of auxiliary domain iron-sulfur clusters is modulated by substrate binding.

On a broader scale, genome-mining studies using the darobactin biosynthetic operon as a template have subsequently uncovered several related biosynthetic gene clusters encoding cyclophane-forming radical SAM enzymes and leading to the discovery of additional bicyclic peptide antibiotics containing N-C and C-C crosslinks.^32,87,88^ The rapid expansion of this enzyme family highlights the importance of understanding how these enzymes recognize their precursor peptides and assemble productive catalytic complexes. The substrate-recognition mechanism identified here therefore provides a new principle for examining related RiPP biosynthesis. Moreover, using an N-terminal extension rather than a canonical RRE domain for precursor-peptide recognition in the presence of SAM expands the known strategies by which RiPP-modifying radical SAM enzymes engage their substrates. Defining these recognition principles will also be important for understanding substrate tolerance and, ultimately, for evaluating the potential of RiPP-forming radical SAM enzymes as platforms for generating modified peptide scaffolds. Furthermore, this study expands the implementation of anaerobic HDX-MS as a useful tool in probing more sensitive or difficult anaerobic systems in the future.

## Acknowledgement

This work is supported by Florida State University (FSU) start-up funds (W.Z.). This work is also in part supported by the National Institutes of Health (R35GM160337 and R03NS147056 to W.Z. and 1S10OD020062-01 to A.T.I.). The content is solely the responsibility of the authors and does not necessarily represent the official views of the National Institutes of Health. The EPR measurement was performed at the University of Wisconsin-Milwaukee. Additional EPR and training support was provided by the National High Magnetic Field Laboratory (NHMFL), supported by National Science Foundation (NSF) Cooperative Agreements DMR1644779 and DMR2128556, and by the State of Florida. Access to the NHMFL EPR instrument was supported by NSF (CHE-2320338 to W.Z.). The DarE and pPH151 plasmids were a gift from Dr. Judith P. Klinman at the University of California, Berkeley. The authors thank Dr. Tomas Orlando at the NHMFL for additional EPR support, Dr. Huan He at the FSU College of Medicine Translational Science Laboratory and Dr. Xinsong Lin at the FSU Department of Chemistry and Biochemistry MASS facility for additional mass spectrometry support.

## Notes

The authors declare no competing financial interests.

## Author Contributions

W.Z. conceived experiments; L.E.C. conducted DarE sample preparation and HDX-MS sample labeling, data curation and analysis; S.K.K. developed methods and prepared recombinant DarA; J.W. performed and analyzed EPR experiments and data analysis; A.T.I. ran the HDX-MS samples; W.Z., L.E.C and J.W. wrote the initial draft, with contributions from all authors.

## TOC

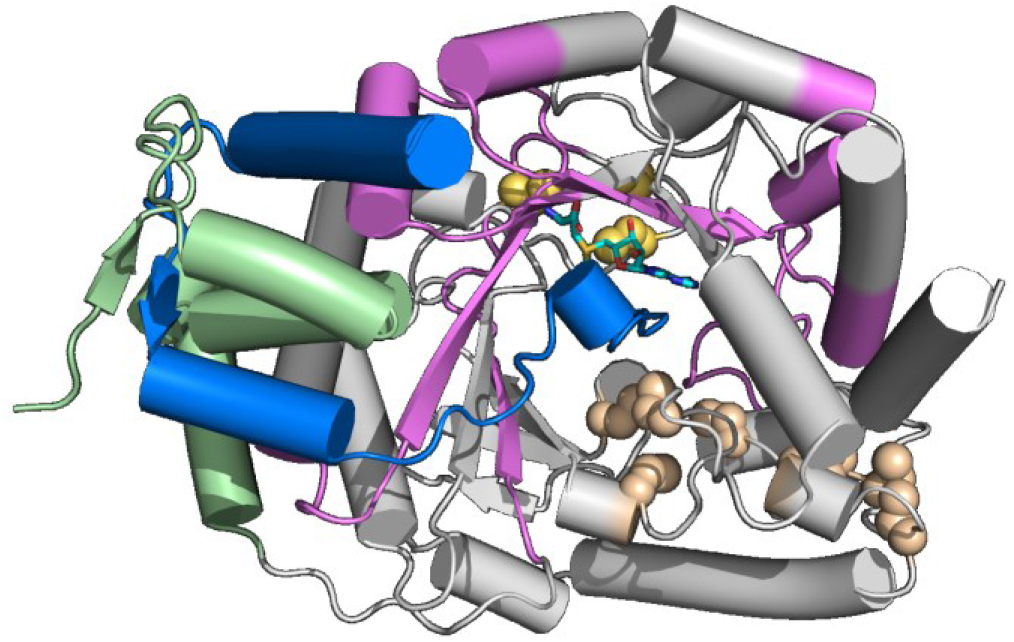

