## Supplementary Information for "SAM-Dependent Precursor Peptide Engagement by Darobactin Maturase DarE"

### Materials and Methods

**General Information.** All chemicals and reagents were purchased from Fisher Scientific or Merck MilliporeSigma unless otherwise stated. Recombinant plasmid DNA constructs, DarA peptide (95% purity), and fluorescein isothiocyanate (FITC)-labeled DarA (95% purity) were synthesized by GenScript (Piscataway, NJ). DNA sequences were confirmed at the Biological Core Facility at Florida State University. Terrific Broth (TB) media was purchased from Research Products International. Anaerobic experiments were conducted in an mBraun Unilab Pro SP anaerobic chamber (Stratham, NH) with $O_2$ levels maintained below 1 ppm. All mass spectrometry analysis of HDX and enzyme activity samples were performed at the University of California, Berkeley QB3 Mass Spectrometry Facility.

**DarE Expression.** The gene encoding *Photorhabdus khanii* DarE (UniProt ID: A0A7C9KC68) with an N-terminal Strep-tag was codon-optimized for *E. coli* expression, synthesized, and inserted into the pET-28b(+) vector between *NcoI* and *XhoI* restriction sites to generate the pET28b-StrepDarE plasmid. Sanger DNA sequencing was performed using T7 forward and reverse primers. Plasmids pET28b-StrepDarE and pPH151, an iron-sulfur assembly helper plasmid, were co-transformed into *E. coli* BL21-Gold (DE3) competent cells and grown overnight in Luria Broth (LB) media containing 50 µg/mL kanamycin and 35 µg/mL chloramphenicol at 37 °C and 220 rpm. For a routine 8 L protein expression, each 1 L of freshly made TB media containing the previously stated antibiotics and 150 µM ammonium iron (III) citrate in a 2 L flask was inoculated with 12 mL of the overnight culture. Cell culture was shaken at 37 °C, 220 rpm until an optical density ($OD_{600}$) of 0.6-0.8 was reached before lowering the temperature and rotation speed to 18 °C and 90 rpm for 1 hr. The protein expression was initiated with 150 µM Isopropyl β-D-1-thiogalactopyranoside (IPTG) and supplemented with 1.5 mM freshly prepared L-cysteine and 750 µM magnesium sulfate. Cells were harvested by centrifugation at 4000 rpm for 20 min at 4 °C after 26 hr of growth and stored in liquid nitrogen until purification.

**DarE Purification.** All purification steps were performed anaerobically using a protocol adapted from previously published work.[1,2] Buffers used for DarE purification were degassed for 1 hr before equilibration under anaerobic conditions overnight. Solutions requiring freshly weighed components added either inside or outside the anaerobic chamber were dissolved in pre-equilibrated anaerobic buffers, followed by extensive purging of the chamber atmosphere during equilibration to maintain anaerobic conditions. Briefly, the cell pellet obtained from an 8 L expression was resuspended with stirring in 30 mL of Buffer A (50 mM EPPS, 150 mM NaCl, pH 8) containing 970 µM DL-dithiothreitol (DTT), 1× BugBuster® protein extraction reagent (Millpore, MA), 6.7 U/mL Benzonase® Nuclease (Millpore, MA) and 0.33 mg/mL lysozyme (Millipore, MA) and incubated for 1 h. The resulting cell lysate was removed from the glovebox in a sealed tube and centrifuged at 12,000 rpm for 45 min at 4 °C before being brought back to the anaerobic chamber under anaerobic conditions for the remainder of the purification. The supernatant was loaded onto a 10 mL Strep-Tactin®XT 4Flow® high-capacity column equilibrated in Buffer A and washed with 30 mL of Buffer A. DarE was eluted with Buffer B (50 mM EPPS, 150 mM NaCl, 2.5 mM D-desthiobiotin, pH 8) and the dark brown colored fractions were collected (Figure S1). The purified sample was buffer-exchanged into Buffer A using a PD-10 column and concentrated with a 0.5 mL 30 kDa Amicon® Ultra Centrifugal Filter Unit at 14,000 × g in 12 min increments. Concentrated DarE was stored in sealed tubes in liquid nitrogen until further use. Protein concentrations were initially determined using the Bradford assay with bovine serum albumin as the standard.[3] To obtain a more accurate estimate of DarE concentration, identical triplicate samples were independently analyzed by amino acid analysis (University of California, Davis, Molecular Structure Facility) following a previously published procedure.[4] Comparison of the concentrations obtained from amino acid analysis and the Bradford assay yielded a correction factor of 0.73, which was subsequently applied to all DarE concentration measurements.

Specifically, concentrations determined by routine Bradford assays were multiplied by this correction factor. Using this procedure, approximately 0.6 mg of DarE was obtained per liter of cell culture expression.

**UV-vis Spectroscopy.** As-purified DarE was diluted under anaerobic conditions to 12.5 µM in the presence or absence of 2 mM DTH, transferred to a 1 mL quartz cuvette, and sealed. The cuvette was then removed from the anaerobic chamber, and the absorbance was recorded using an Agilent Cary 60 UV-vis spectrophotometer (Figure S1).

**Iron Content Quantification.** The iron content of DarE was determined by following previously established protocols.[5,6] A range of iron standards from 0-1 mM was prepared in MilliQ water to generate a calibration curve. A 100 µL aliquot of each iron standard solution and triplicate DarE samples were incubated with 100 µL of 22% $HNO_3$ at 90 °C for 30 min. After cooling to room temperature, the samples were centrifuged at 12,000 × g for 1 min, and the supernatant was neutralized by adding 600 µL of 7.5% ammonium acetate before adding 100 µL 12.5% freshly made ascorbic acid and 100 µL of 10 mM ferrozine reagent. The resulting solutions were centrifuged again at 12,000 × g for 1 min before transferring to cuvettes, and the absorbance at 562 nm was recorded using an Agilent Cary 60 UV-vis spectrophotometer. The iron content of DarE was calculated using the linear regression equation obtained from the iron-standard calibration curve.

**Sulfide Content Quantification.** The sulfur content of DarE was determined using the previously reported methods under anaerobic conditions.[5,6] A range of sodium sulfide standards from 0-1000 µM was prepared in MilliQ water to generate a calibration curve. A 200 µL aliquot of each sodium sulfide standard and a triplicate of DarE samples were incubated for 15 min at room temperature with 600 µL 1% (w/v) freshly made zinc acetate and 50 µL 12% (v/v) NaOH. The samples were then centrifuged at 14,000 × g for 1 min before adding 150 µL of 0.1% freshly made *N,N*-dimethyl-*p*-phenylenediamine dihydrochloride in 5 M HCl and 150 µL of freshly made 10 mM $FeCl_3$ in 1 M HCl. The solutions were mixed by inversion and incubated for 30 min before being removed from the anaerobic chamber and transferred to cuvettes, and the absorbance was read at 670 nm using an Agilent Cary 60 UV-vis spectrophotometer. The sulfide content per monomer of DarE was calculated using the linear regression equation obtained from the sulfide-standard calibration curve.

**Size-exclusion Chromatography**. Various dilutions of DarE, ranging from 10-100 µM, were prepared under anaerobic conditions, centrifuged at 12,000 × g for 10 min, and injected into a Cytiva Superdex 200 Increase size-exclusion chromatography column that was previously equilibrated with degassed Buffer A. The absorbance at 280 nm was measured as the column was eluted with Buffer A at a 0.22 mL/min flow rate at 4 °C. To determine the molecular weight of DarE corresponding to each peak, the retention times were compared to a standard curve generated from the commercially available Sigma Protein Standard Mix (15-600 kDa) that was measured under the same conditions. Peak areas were integrated using Cytiva UNICORN 7.10 evaluation software, and the fractional peak area for both monomer and dimer were calculated by dividing the peak area of each by the total peak area of both monomer and dimer combined at each DarE concentration. Then, the concentration of monomer and dimer was determined in each sample by multiplying the fractional amount of each by the total DarE concentration. Using the determined monomer and dimer concentrations, the monomer-dimer equilibrium expression was applied to calculate the approximate $K_d$, and the average between conditions was reported.

**DarA Expression:** Besides the synthetic DarA, we have also developed a recombinant *Photorhabdus khanii* DarA preparation protocol. DarA (UniProt ID: W3V7B0) was expressed as a $His_6$-SUMO-tagged fusion protein (Table S4). Briefly, the N-terminal SUMO-fused DarA sequence was codon-optimized, flanked with 5'-*NdeI* and 3'-*BamHI* restriction sites, and inserted into the pET-28b(+) vector. The sequence of insertion was confirmed by Sanger sequencing and transformed into *E. coli* BL21-Gold (DE3) competent cells. Cells containing the SUMO-DarA plasmid were grown overnight in 10 mL LB media supplemented with 50 µg/mL kanamycin at 37 °C at 200 rpm. The overnight culture was used to inoculate 1 L of LB media supplemented with 50 µg/mL of kanamycin in a 2.4 L Fernbach flask. The flasks were then placed in a shaking incubator set at 37 °C until the $OD_{600}$ at 600 nm approximately reached 0.6-0.8. The incubator temperature was further lowered to 18°C, and the flasks were incubated for an additional hour. IPTG was added to induce protein expression at a final concentration of 0.5 mM as the flasks were shaken for an additional 18 hr. The cells were harvested by centrifugation at 3,000 × g at 4 °C for 30 min and the pelleted cells were resuspended in Buffer C (20 mM Tris-Cl, 300 mM NaCl, and 10% *v/v* glycerol, pH 7.5). The resuspended cells were then frozen and stored at -80 °C until purification.

**DarA Purification** Cells were thawed and lysed by sonication on ice using a Qsonica Q500 sonicator, with a standard tip at 30% amplitude for 6 min with 15 s ON and 45 s OFF. The lysate was clarified by centrifugation at 15,000 × g for 30 min to remove insoluble cellular debris. Protein purification was performed at room temperature using a 2 mL Ni-NTA column. All buffers were filtered through 0.22 µm polyvinylidene difluoride membrane filters. The supernatant was loaded onto a column equilibrated with Buffer D (20 mM Tris-Cl, 1 M NaCl, 30 mM imidazole, pH 7.5). After loading the sample, the column was washed with 5 column volumes of Buffer D. The $His_6$-tagged protein was then eluted with 5 column volumes of Buffer E (20 mM Tris-Cl, 1 M NaCl, 250 mM imidazole, pH 7.5). The purity of the eluted fractions was assessed using SDS-PAGE, and fractions containing pure protein were pooled together (Figure S12). The pooled fractions were desalted by repeated passage through a pre-equilibrated Sephadex™ G-25 PD-10 desalting column (Cytiva) equilibrated with 30 mL of Buffer F (20 mM Tris-Cl, 150 mM NaCl, 10 mM imidazole, pH 7.5), loading 2.5 mL of sample and eluting with 3.5 mL of Buffer F per pass, repeated until the entire pooled volume had been processed. The eluted fractions were pooled, and $His_6$-tagged ULP1 protease was added, followed by incubation at room temperature for 4 hr. The extent of SUMO cleavage was determined with an SDS-PAGE using full-length SUMO-DarA as a control. The cleaved protein was loaded onto a 2 mL Ni-NTA column equilibrated with Buffer F, and the flow-through was collected. The column was washed with 2 column volumes of Buffer F and pooled with the flow-through previously collected. The pooled fractions were acidified with formic acid to a final concentration of 0.1% (v/v) and loaded onto a HyperSep™ C18 SPE column (500 mg/6 mL, Thermo Scientific) pre-washed with acetonitrile containing 0.1% v/v formic acid (FA), followed by water with 0.1% v/v FA. The column was subsequently washed with 0:100, 75:25, and 100:0 mixtures of acetonitrile (0.1% v/v FA):water (0.1% v/v FA). The presence of cleaved peptide was verified by measuring A280 on a NanoDrop-2000c (Thermo Scientific) for the 75:25 and 100:0 fractions. The peptide-containing fractions were pooled and dried in a Lyophilizer (Labconco). The dried peptide was resuspended in water to a target concentration of 2.5 mM, calculated from A280 using a molar extinction coefficient of 11,000 $M^{-1}cm^{-1}$,[7] and the remaining dried powder was stored at -80 °C until further use.

**Flavodoxin and Flavodoxin-NADPH Reductase Protein Preparation** The gene encoding *E. coli* flavodoxin (*Ec*FldA, UniProt ID: P61949) and flavodoxin $NADP^+$ reductase (*Ec*FNR, UniProt ID: P28861) with N-terminal His-tags were synthesized and inserted into the pET-28b (+) vector between *NdeI* and *BamHI* restriction sites. Plasmids containing *Ec*FldA or *Ec*FNR genes were transformed into *E. coli* BL21-Gold (DE3) competent cells and grown overnight in LB media containing 50 µg/mL kanamycin at 37 °C and 220 rpm. For protein expression, each one liter of LB media was inoculated with 10 mL of overnight culture and grown to an $OD_{600}$ of 0.7. Protein expression was induced by adding 300 µM IPTG and 17.5 µM flavin mononucleotide (FMN), and the temperature and rotation were lowered to 30 °C and 200 rpm, respectively. Cells were harvested at 4000 rpm at 4 °C for 20 min after 16 hr of expression and stored at -80 °C until purification. *Ec*FldA and *Ec*FNR were purified following the same protocol adapted from the previous publication.[8] The cell pellet was resuspended in 40 mL Buffer G (50 mM EPPS, 150 mM NaCl, 30 mM imidazole, pH 8), sonicated at 25% amplification, 1 s ON and 1 s OFF for 6 min. For *Ec*FNR purification, 40 μM flavin adenine dinucleotide (FAD) was added to the cell lysate during sonication. The lysate was then centrifuged at 12,000 rpm for 30 min at 4 °C. The supernatant was loaded onto a 6 mL Ni-NTA column equilibrated with Buffer G. The column was washed with Buffer G and eluted with Buffer H (50 mM EPPS, 150 mM NaCl, 300 mM imidazole, pH 8) while collecting fractions. Elution fractions containing protein were then combined and concentrated using an Amicon® Ultra 10 kDa cutoff centrifugal filter. After buffer exchange to Buffer A using a PD-10 column, the protein was concentrated again and stored at -80 °C until further use. Protein concentration was determined using a Bradford assay. As-purified protein samples were then scanned from 200-900 nm using an Agilent Cary 60 UV-vis spectrophotometer to confirm cofactor content (Figure S12). Using the estimated extinction coefficients of the absorbances at 280 and 450 nm corresponding to the protein and cofactor, respectively, the concentrations of each were determined. The ratio of FMN:*Ec*FldA was 0.114 and of FAD:*Ec*FNR was 0.355. The protein concentration was corrected for cofactor loading.

**Electron Paramagnetic Resonance (EPR) Spectroscopy.** Samples for X-band (~9.4 GHz) EPR spectroscopy were prepared under anoxic conditions (<0.5 ppm $O_2$) in an LC-100 glovebox (LC Technology Solutions Inc.) (Figure S8). Oxidized samples were prepared by freezing approximately 250 μL of 100-200 μM DarE. Reduced samples were prepared by mixing approximately 225 μL of 100-200 μM DarE with 25 μL of a 100 mM stock of dithionite for a final concentration of 10 mM followed by a short incubation (1-2 minutes). Samples containing 10 mM *S*-adenosyl L-methionine (SAM) and/or 888 μM DarA were incubated before reduction to a final volume of 225 μL before adding dithionite as above. The prepared samples were then added to a 4 mm OD quartz sample tube (Wilmad) and immediately frozen in the glovebox in an ethanol/dry ice bath followed by removal from the glovebox while frozen and stored in liquid nitrogen until use. Continuous wave (CW) spectra were collected using a Magnettech MS5000 spectrometer (Freiberg Instruments) equipped with a Cold-Edge (Allentown, PA) Scorpio-TT cryostat and Stinger closed loop helium cooling system. Sample temperatures were monitored and maintained using a Lake Shore Cryotronics 336 temperature controller. Spectra were recorded at 10 K using 9.4 GHz microwave frequency, 2 mW microwave power, 120 s sweep time 0.5 mT modulation amplitude. Spectra analysis, plotting, and simulations were performed using Easyspin (Easyspin.org) in MATLAB R2019a (Mathworks Inc). Spectra are shifted from their collection frequencies, which vary from 9.465 to 9.485 GHz, to a common 9.475 GHz for plotting and spectral comparison by using the following equation:

$$\mathrm{x(data\ @9.475GHz) = x(data\ @\ orig\ freq.) * \frac{9.475}{\nu(of\ collected\ data\ in\ GHz)}}$$

**DarE Activity Assay and LC-MS Analysis.** $O_2$-containing buffer was prepared by saturating Buffer A with air for 90 min at 1 atm prior to sample preparation. The $O_2$ concentration in this air-saturated buffer solution is estimated to be 260 μM.[9] An aliquot of 50 μM DarE was incubated with 300 μM DarA and 1 mM SAM for 15 min under anaerobic conditions before the addition of 25 μM *Ec*FldA, 2 μM *Ec*FNR, 10 mM NADPH, and ~75 μM $O_2$-containing Buffer A. After 0 min-18 hr of incubation at room temperature, the reaction was quenched with 2 μL formic acid and centrifuged at 12,000 rpm for 10 min. Supernatant was analyzed using high-resolution LC-MS at the University of California, Berkeley QB3 Mass Spectrometry Facility using the methods established previously.[10] The time course of the activity curve was fitted to a single exponential: $[5'\text{-dAdo}](t) = Y_0 + (\text{plateau}-Y_0)(1-e^{-kt})$.

**Fluorescence Polarization Assay.** The interaction between DarA and DarE was determined using fluorescence polarization in a competitive assay format in a 10 μL sample size. Briefly, 1 nM DarE was incubated with 1 nM N-terminal FITC-labeled DarA for 10 min. A serial dilution of unlabeled DarA peptide solution was titrated into the mixture in a UV-STAR® small volume μCLEAR® black 384-well microplate in the anaerobic chamber. The plate was sealed with polyester adhesive film before being centrifuged at 1000 rpm for 1 min. The fluorescence polarization of samples was determined using a BMG Labtech VantaStar™ plate reader. FITC was excited at 482 nm, and emission was recorded at 530 nm. The fluorescence polarization readings were plotted versus DarA concentration, and the data was fit using GraphPad Prism. For the sample containing SAM, the same experiment was performed in the presence of 2 mM SAM. For control, DarA was titrated into the mixture containing the probe and buffer and omitted DarE. The measurements with and without SAM were performed on different days, and DarE was present far below the apparent $IC_{50}$ values. Both synthetic DarA and recombinant DarA were tested in triplicated experiments. Although batch-to-batch variation was observed in synthetic DarA (GenScript, NJ), the SAM-dependent apparent $IC_{50}$ shift was reproducible across various DarE and recombinant DarA.

**Peptide Library Establishment Using LC-MS/MS.** To optimize and evaluate proteolysis for DarE, stock solutions of DarE and DarA were placed in an anaerobic chamber and equilibrated for 30 min. Master mixtures of protein samples containing 25 μM DarE and 50 μM DarA, with or without 5 mM SAM in Buffer A, were prepared. After 15 min of incubation, 10 μL of master mixture was diluted to 50 μL in $H_2O$-based Buffer A to obtain a labeling concentration of 5 μM DarE and 10 μM DarA, with or without 1 mM SAM. Reactions were quenched with 15 uL of Quenching Buffer (2 M citric acid, 3 M guanidine-HCl, 3 mM TCEP, pH 1) at -4 °C for 2 min. Proteolysis was initiated by mixing 6.5 μL of 2 mg/mL pepsin and 3.5 μL of 2 mg/mL protease from *Aspergillus saitoi* Type XIII and incubating on ice for 13 min. A 60 μL aliquot of each sample was transferred to an MS analysis tube, flash-frozen, and stored in liquid nitrogen before MS analysis. The LC-MS/MS analysis protocol was adapted from the previous publication.[10]

**HDX Sample Preparation and Measurements** All H/D exchange samples were prepared in the anaerobic chamber. Stock solutions of DarE and DarA stored outside the anaerobic chamber were equilibrated under anaerobic conditions for at least 30 min before setting up the labeling reactions. Master mixtures of protein samples were prepared for 1) 25 μM DarE, 2) 25 μM DarE + 50 μM DarA, 3) 25 μM DarE + 5 mM SAM, and 4) 25 μM DarE + 5 mM SAM + 50 μM DarA in Buffer A. A 10 μL aliquot of master mixture was incubated for 15 min before being diluted to 50 μL in $D_2O$-based Labeling Buffer (50 mM EPPS, 10 mM TCEP, 30 mM KCl, pD 8) so that the final concentrations during labeling were 5 μM DarE, 10 μM DarA, and/or 1 mM SAM. Labeling reactions were quenched with 15 μL of Quenching Buffer (2 M citric acid, 3 M guanidine-HCl, 3 mM TCEP, pH 1) at 45 s, 600 s, 1800 s, 3600 s, and 10800 s at -4 °C for 2 min. Proteolysis was initiated by mixing 6.5 μL of 2 mg/mL pepsin (ThermoFisher Cat. J61679.06) and 3.5 μL of 2 mg/mL protease from *Aspergillus saitoi* Type XIII (SigmaAldrich Cat. P2143-5G) and incubating on ice for 13 min. A 60 μL aliquot of each sample was transferred to an MS analysis tube, flash-frozen, and stored in liquid nitrogen prior to MS analysis. The 0 s labeling samples were prepared using the same procedure as the 2 min time point, except that $H_2O$-based Buffer A was used. All time points for each condition were prepared on the same day, and the protocol was repeated on a different day for the duplicate samples. Labeled samples were stored in liquid nitrogen for less than a week before analysis by high-resolution LC-MS. The LC-MS method was adapted from the previous publication.[10]

**HDX-MS Data Analysis.** The resulting raw MS data were analyzed using HDX WorkBench[11], and the spectra of detected peptides were manually curated to ensure accurate detection and analysis of exchange behavior. A set of high-quality peptides derived from DarE and DarA across conditions was selected for further analysis. The H/D exchange data, expressed as percent deuterium uptake (%D) at each time point for each peptide, were plotted using PRISM GraphPad. For each peptide, the cumulative %ΔD across all five labeling time points was calculated and plotted. A Σ$\Delta$%D cutoff value of 25 was applied to define notable differences between the apo protein and the corresponding binary and ternary complexes, consistent with the standard criteria used in previous HDX-MS studies.[12–14]

**OpenFold3 analysis.** To predict the binding of DarA to DarE, corresponding sequences were input and processed using online access to OpenFold3 through the Neurosnap webserver. SAM was input through the built-in database search feature, which referenced the PubChem CID (34755). The resulting output for the analysis was five separate models, all ranked with a “very high” overall quality and aggregate score above 0.9, indicating high confidence. A full summary of the analysis output is provided in Table S4. Each model ranking was visualized and compared via PyMOL (Figures S7 & S9).

**Table S1. PDB code and conformational analysis reported for the SPASM domain-containing RiPP-producing radical SAM enzymes.**

| Enzyme | Apo | SAM/SAH-bound | Peptide-bound | Conformational change | Reference |
|---|---|---|---|---|---|
| SuiB | 5V1Q | 5V1S | 5V1T (with SAM) | Yes | [27] |
| PqqE | 6C8V | - | - | Yes (HDX-MS) | [10,28] |
| SkfB | - | 6EFN | - | - | [29] |
| EpeE | - | 8AI1 | 8AI2, 8AI5, 8AI4 (with SAH) | No | [30] |
| TigE | 8VPO | - | - | - | [31] |
| CteB | - | 5WHY | 5WGG (with SAM) | No | [32] |

**Table S2. The *g*-tensors reported for other SPASM domain-containing radical SAM enzymes.**

| Enzyme | Species | *g*-values | *g*-avg | Reference |
|---|---|---|---|---|
| DarE | $[4Fe\text{-}4S]^{+}_{RS+SAM}$ | [1.98836, 1.89177, 1.833] | 1.9044 | This work |
| | $[4Fe\text{-}4S]^{+}_{ClusterA}$ | [2.045, 1.920, 1.850] | 1.9383 | |
| | $[4Fe\text{-}4S]^{+}_{ClusterB}$ | [1.9751, 1.9841, 1.9147] | 1.9580 | |
| PqqE | $[4Fe\text{-}4S]^{+}_{RS}$ | [2.040,1.927,1.897] | 1.9547 | 15 |
| | $[2Fe\text{-}2S]^{+}_{AuxI}$ | [2.004,1.958,1.904] | 1.9553 | |
| | $[4Fe\text{-}4S]^{+}_{AuxII}$ | [2.059,1.940,1.903] | 1.9673 | |
| SuiB | $[4Fe\text{-}4S]^{+}_{RS}$ | [2.048,1.936,1.916] | 1.9667 | 16 |
| | $[4Fe\text{-}4S]^{+}_{AuxI}$ | [2.060,1.939,1.904] | 1.9677 | |
| | $[4Fe\text{-}4S]^{+}_{AuxII}$ | [2.057,1.946,1.872] | 1.9583 | |
| Tte1186 | $[4Fe\text{-}4S]^{+}_{RS}$ | [2.04,1.92,1.91] | 1.9567 | 17 |
| | $[4Fe\text{-}4S]^{+}_{RS+SAM}$ | [1.99,1.90,1.84] | 1.9100 | |
| | $[4Fe\text{-}4S]^{+}_{AuxI}$ | [2.06,1.93,1.88] | 1.9567 | |
| | $[4Fe\text{-}4S]^{+}_{AuxII}$ | [2.05,1.93,1.89] | 1.9567 | |
| RumC1 | $[4Fe\text{-}4S]^{+}_{RS}$ | [2.034,1.913,1.870] | 1.9390 | 18 |
| | $[4Fe\text{-}4S]^{+}_{RS+SAM}$ | [2.076,1.930,1.852] | 1.9527 | |
| | $[4Fe\text{-}4S]^{+}_{ClusterA}$ | [2.050,1.937,1.902] | 1.9630 | |
| | $[4Fe\text{-}4S]^{+}_{ClusterB}$ | [2.074,1.932,1.885] | 1.9637 | |
| AlbA | $[4Fe\text{-}4S]^{+}_{RS}$ | [2.03,1.92] | 1.9567 | 19 |
| | $[4Fe\text{-}4S]^{+}_{ClusterA}$ | [1.92,2.03] | 1.9567 | |
| GggB | $[4Fe\text{-}4S]^{+}_{RS}$ | [2.03,1.91,1.88] | 1.9400 | 20 |
| NxxcB | $[4Fe\text{-}4S]^{+}_{RS}$ | [2.040,1.905,1.900] | 1.9483 | 21 |
| BtrN | $[4Fe\text{-}4S]^{+}_{RS}$ | [2.04,1.92] | 1.9600 | 22 |
| | $[4Fe\text{-}4S]^{+}_{RS+SAM}$ | Peak broadening | | |
| MoaA | $[4Fe\text{-}4S]^{+}_{Cluster\ I}$ | [1.900,2.028] | 1.9427 | 23 |
| | $[4Fe\text{-}4S]^{+}_{Cluster\ II}$ | [1.879,2.060] | 1.9393 | |
| MftC | $[4Fe\text{-}4S]^{+}_{RS}$ | [1.88,1.90,2.05] | 1.9433 | 24 |
| | $[4Fe\text{-}4S]^{+}_{RS+SAM}$ | [1.89,1.90,2.015] | 1.9350 | |
| | $[4Fe\text{-}4S]^{+}_{AuxI}$ | [1.90,1.91,2.04] | 1.9500 | |
| | $[4Fe\text{-}4S]^{+}_{AuxII}$ | [1.87,1.92,2.04] | 1.9433 | |
| TYW1 | $[4Fe\text{-}4S]^{+}$ | [2.02, 1.90, 1.86] | 1.9267 | 25 |
| | $[4Fe\text{-}4S]^{+}_{RS+SAM}$ | [1.98, 1.86, 1.83] | 1.8900 | |
| anSME | $[4Fe\text{-}4S]^{+}$ | [2.05, 1.92] | 1.9633 | 26 |
| | $[4Fe\text{-}4S]^{+}_{RS+SAM}$ | [1.98, 1.90, 1.84] | 1.9067 | |

**Table S3. Results output for OpenFold3 analysis for each model ranking of DarE in complex with DarA and SAM.**[33,34]

| Model ID | 1 | 2 | 3 | 4 | 5 |
|---|---|---|---|---|---|
| Overall Quality | Very High | Very High | Very High | Very High | Very High |
| Aggregate Score | 0.947 | 0.936 | 0.927 | 0.927 | 0.92 |
| Avg pLDDT | 82.49 | 82.24 | 82.48 | 82.87 | 82.58 |
| Global PDE | 0.531 | 0.561 | 0.578 | 0.58 | 0.589 |
| pTM | 0.887 | 0.88 | 0.873 | 0.873 | 0.87 |
| ipTM | 0.87 | 0.858 | 0.849 | 0.848 | 0.842 |
| Disorder Fraction | 0.146 | 0.148 | 0.148 | 0.148 | 0.146 |
| Has Clash | No | No | No | No | No |
| Per Chain pTM | DarA: 0.625<br>DarE: 0.881<br>SAM: 0.452 | DarA: 0.610<br>DarE: 0.873<br>SAM: 0.449 | DarA: 0.597<br>DarE: 0.866<br>SAM: 0.444 | DarA: 0.594<br>DarE: 0.866<br>SAM: 0.441 | DarA: 0.585<br>DarE: 0.864<br>SAM: 0.441 |
| Chain Pair ipTM | AE: 0.869<br>AS: 0.569<br>ES: 0.921 | AE: 0.859<br>AS: 0.566<br>ES: 0.920 | AE: 0.849<br>AS: 0.548<br>ES: 0.915 | AE: 0.850<br>AS: 0.549<br>ES: 0.914 | AE: 0.842<br>AS: 0.543<br>ES: 0.914 |
| Bespoke ipTM | AE: 0.807<br>AS: 0.745<br>ES: 0.745 | AE: 0.801<br>AS: 0.743<br>ES: 0.743 | AE: 0.790<br>AS: 0.732<br>ES: 0.732 | AE: 0.791<br>AS: 0.732<br>ES: 0.732 | AE: 0.785<br>AS: 0.729<br>ES: 0.729 |

**Table S4. DNA and protein sequence of recombinant DarA.**

| | **Sequences** |
|---|---|
| SUMO-DarA DNA sequence | AGTGATTCTGAAGTAAATCAAGAGGCCAAGCCGGAGGTGAAGCCAGAGGTCAAGCCGGAAACCCACATCAACCTGAAAGTTAGCGATGGCTCTAGCGAAATTTTCTTCAAAATCAAAAAGACGACCCCGCTGCGCCGTTTGATGGAGGCGTTTGCAAAGCGCCAGGGTAAAGAAATGGACAGCCTGCGTTTTTTGTACGACGGCATTCGTATTCAAGCGGATCAGGCTCCGGAGGACCTGGATATGGAGGATAATGACATCATCGAAGCGCATCGTGAACAGATTGGTGGTATGCACAACACCCTGAACGAAACCGTGAAAACACAAGAGGCACTGAACAGCCTTGCCGCCTCTTTCAAGGAAACGGAACTGTCAATTACCGATAAAGCACTTAACGAACTGAGCAATAAGCCAAAAATCCCTGAAATTACCGCATGGAACTGGTCCAAGTCTTTCCAAGAGATTTAA |
| As-purified SUMO-DarA protein sequence | MGSSHHHHHHSSGLVPRGSHMSDSEVNQEAKPEVKPEVKPETHINLKVSDGSSEIFFKIKKTTPLRRLMEAFAKRQGKEMDSLRFLYDGIRIQADQAPEDLDMEDNDIIEAHREQIGGMHNTLNETVKTQEALNSLAASFKETELSITDKALNELSNKPKIPEITAWNWSKSFQEI |
| Purified DarA peptide sequence | MHNTLNETVKTQEALNSLAASFKETELSITDKALNELSNKPKIPEITAWNWSKSFQEI |

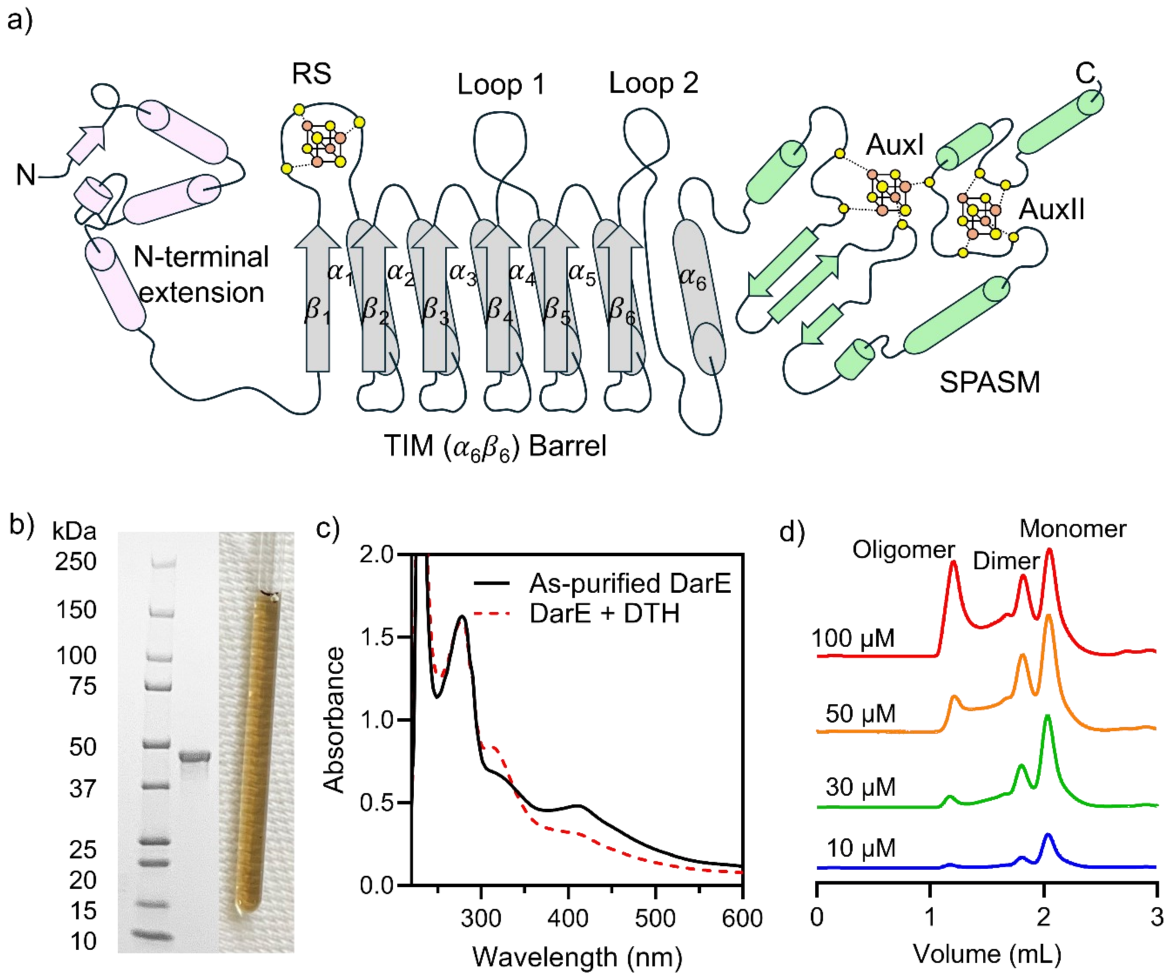


**Figure S1. Predicted DarE secondary structure and sample preparation.** a) Schematic representation of the predicted secondary structure of DarE. b) SDS-PAGE of purified DarE. c) UV-Vis spectrum of as-purified DarE in the absence and presence of sodium dithionite (DTH). D) Size exclusion chromatography profiles of DarE titration.

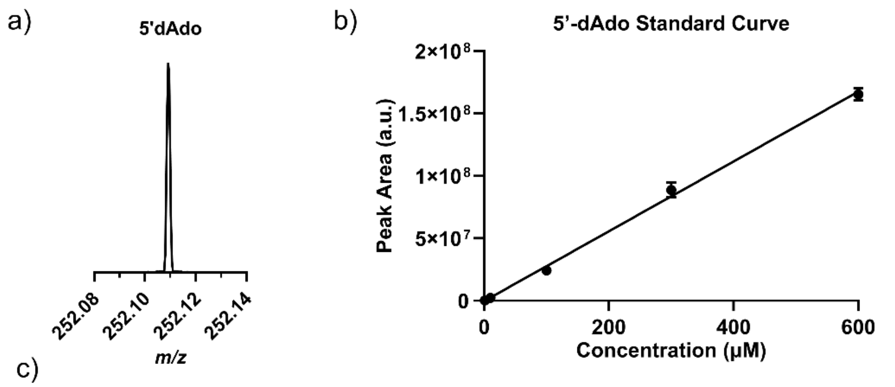


c)

| Time (min) | Peak Area Rep 1 | Peak Area Rep 2 | Rep 1 [5’-dAdo] (μM) | Rep 2 [5’-dAdo] (μM) |
|---|---|---|---|---|
| 1 | 0 | 0 | 0 | 0 |
| 60 | 2.03E+06 | 7.24E+05 | 8.498 | 3.826 |
| 240 | 4.38E+06 | 6.27E+06 | 16.87 | 23.64 |
| 480 | 7.15E+06 | 8.51E+06 | 26.792 | 31.658 |
| 1080 | 1.29E+07 | 1.18E+07 | 47.365 | 43.47 |

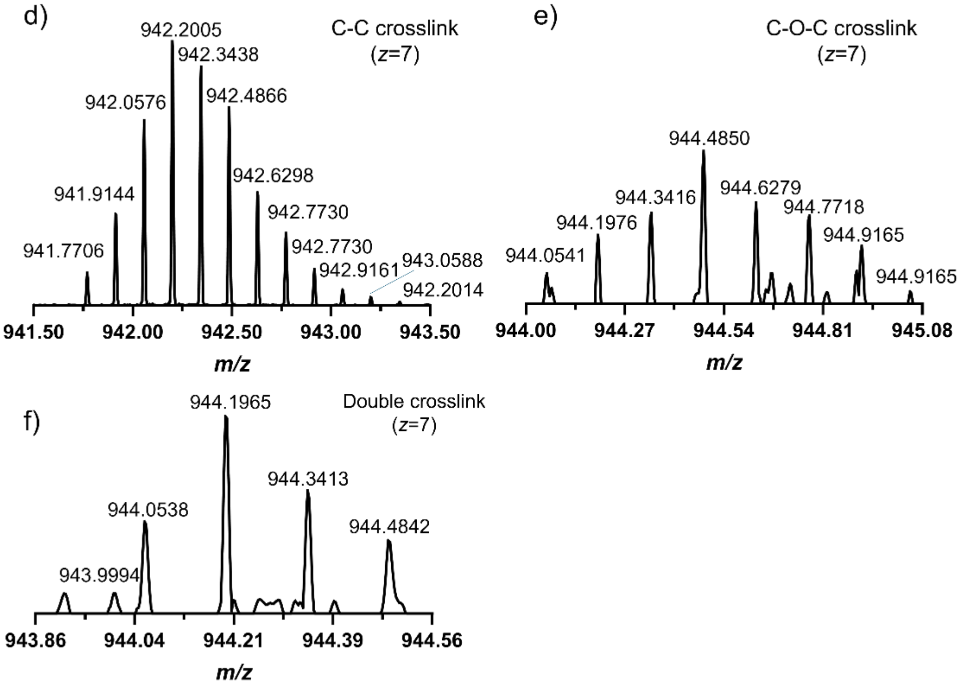


**Figure S2. DarE activity measured by high-resolution mass spectrometry.** a) Extracted ion chromatogram showing the 5’-deoxyadenosine (5’-dAdo) signal. b) Commercially available 5’-dAdo was used to establish a standard curve for quantification of 5’-dAdo production in the DarE-catalyzed reaction. c) Time-dependent formation of 5’-dAdo reported in Figure 2a. with two independent replicates. Peak area was converted to concentration using the standard curve in panel b). d) Representative high-resolution mass spectrometry spectra of modified DarA peptide C-C crosslinks. e) Representative high-resolution mass spectrometry spectra of modified DarA peptide C-O-C crosslinks. f) Representative high-resolution mass spectrometry spectra of modified DarA peptide C-C and C-O-C double crosslinks.

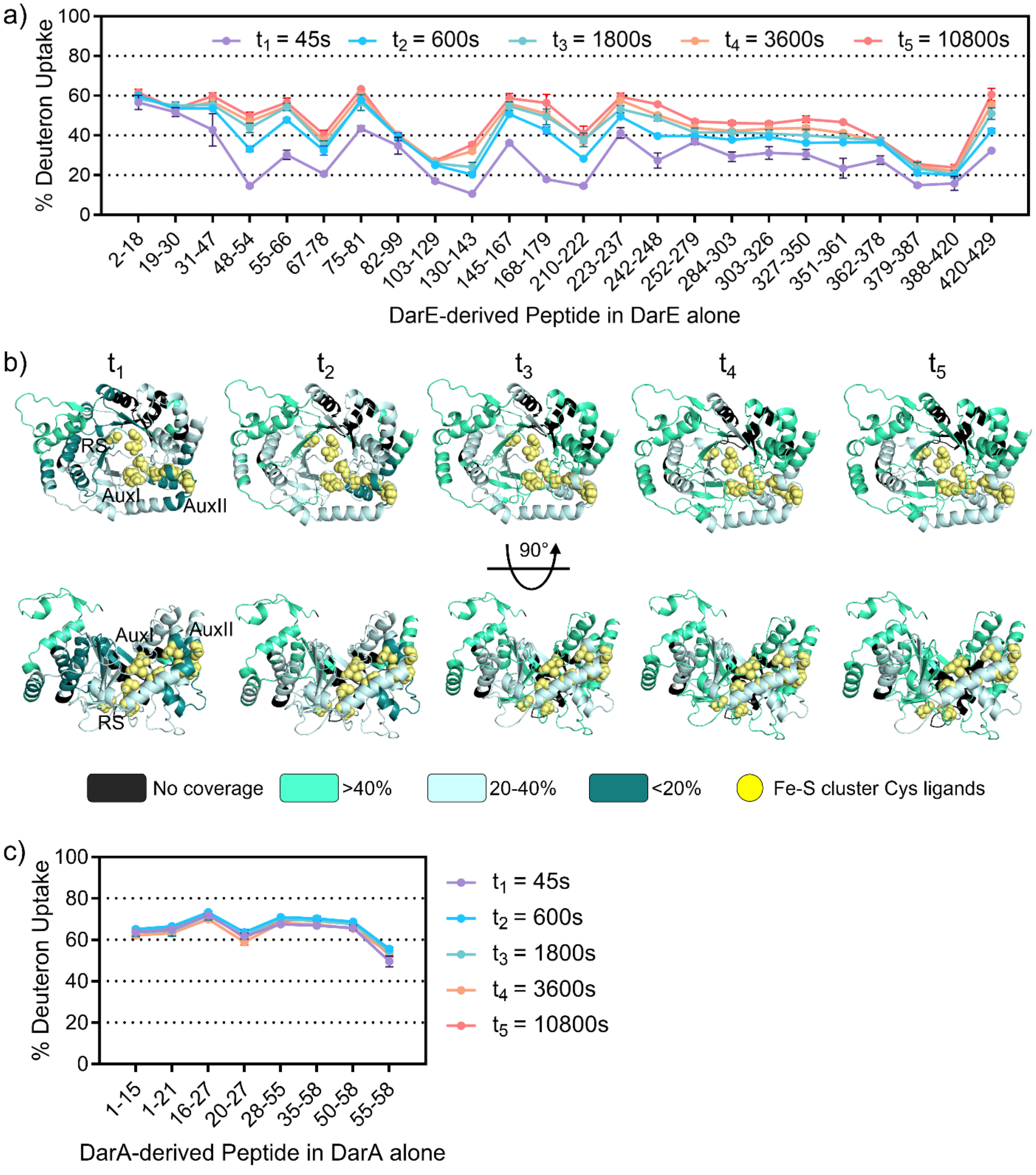


**Figure S3. HDX-MS analysis of DarE and DarA.** a) Deuterium uptake profiles of DarE peptides analyzed in this work across the indicated labeling time point for DarE alone. Data are shown as mean ± sd. b) HDX-MS protection mapped onto the DarE AlphaFold structural model. Regions exhibiting altered deuterium uptake are color-coded as indicated. c) Deuterium uptake profiles of DarA peptides analyzed in this work across the indicated labeling time point for DarA alone. Data are shown as mean ± sd.

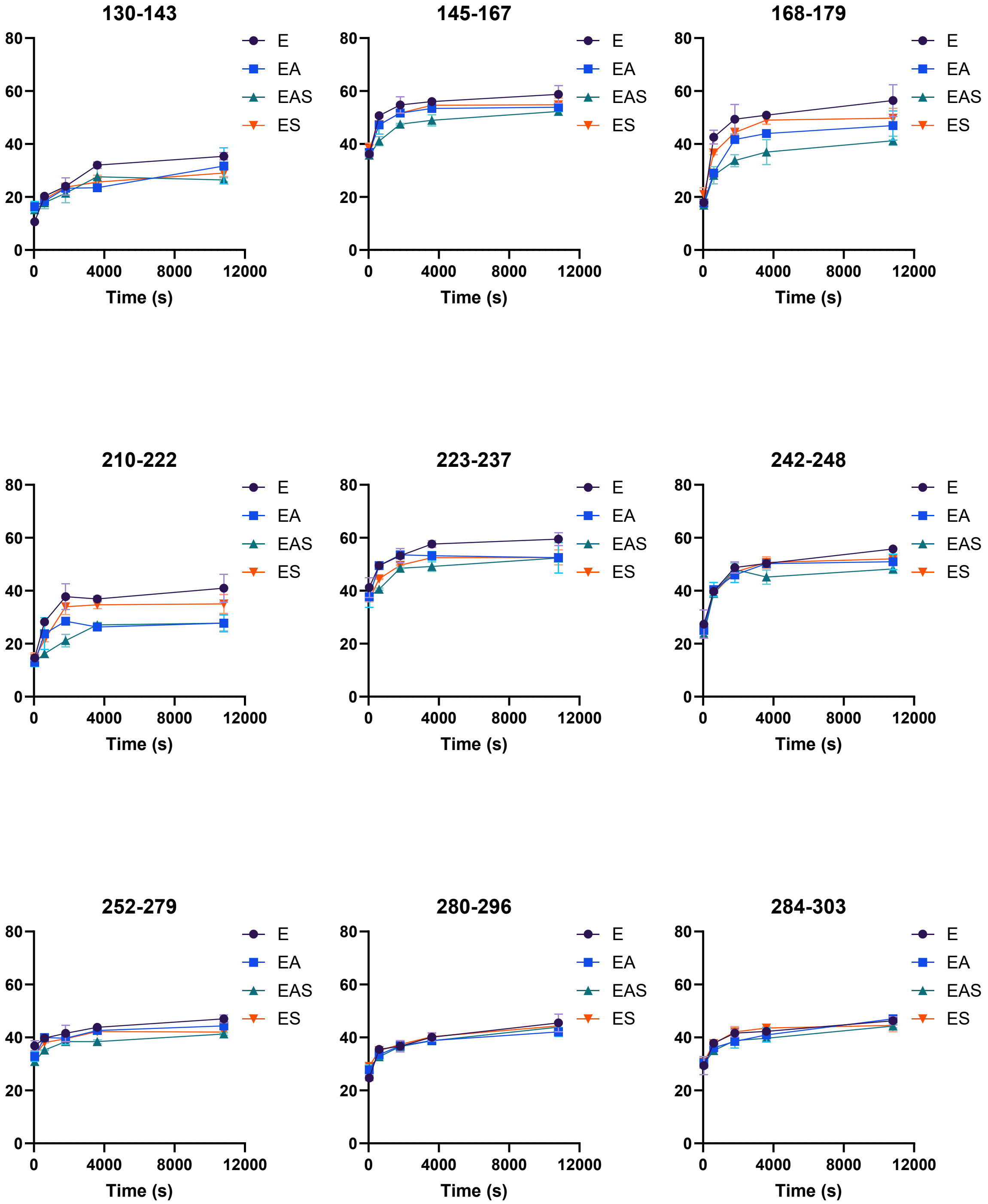


**Figure S4. Percentage deuterium uptake plots of DarE-derived peptides.** Deuterium uptake kinetics are shown for DarE-derived peptides. Percent deuterium uptake was measured for DarE alone (E), DarE-DarA (EA), DarE-DarA-SAM (EAS), and DarE-SAM (ES) across the indicated labeling times. Data are shown as mean ± sd.

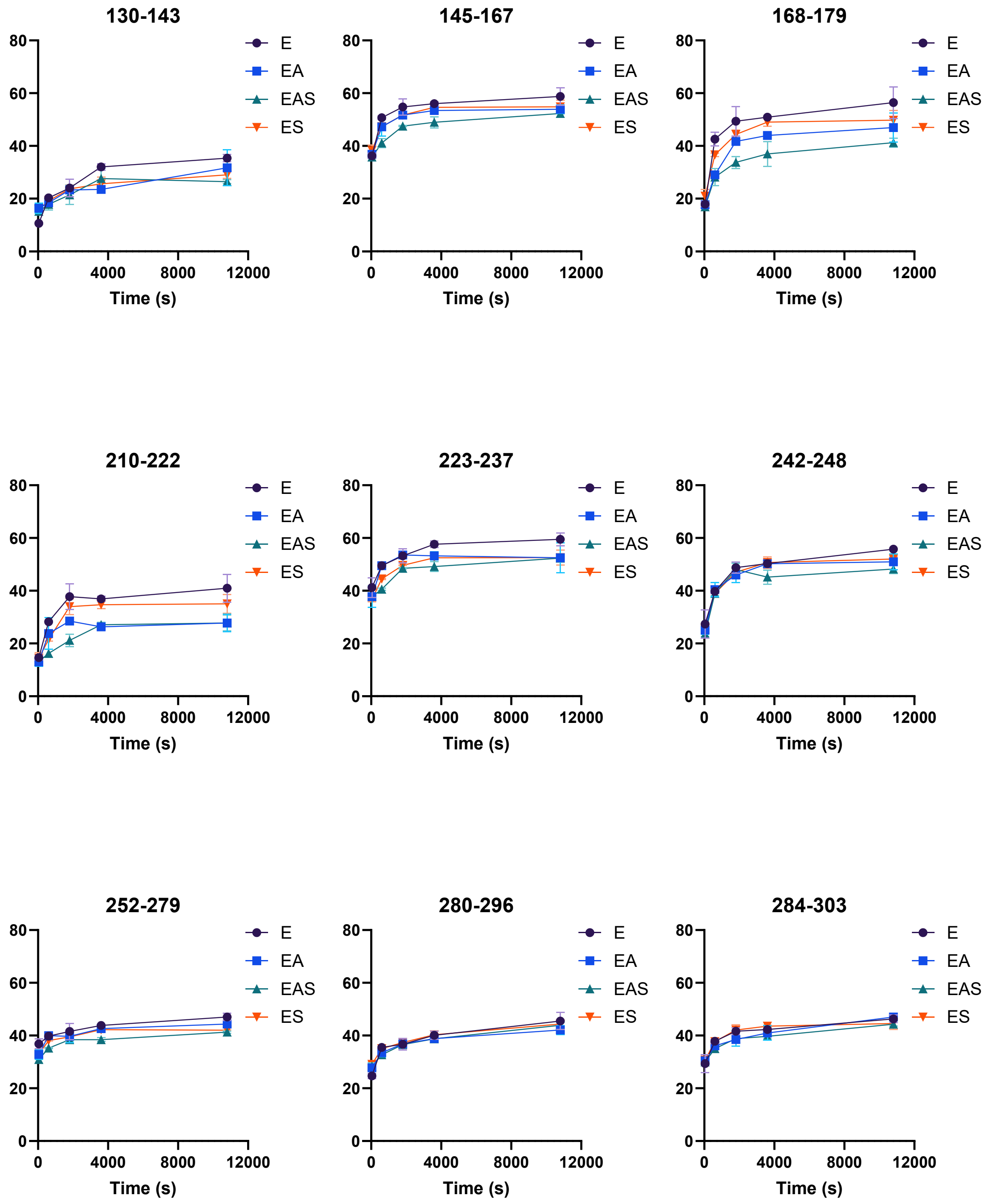


**Figure S4 (continued). Percentage deuterium uptake plots of DarE-derived peptides.** Deuterium uptake kinetics are shown for DarE-derived peptides. Percent deuterium uptake was measured for DarE alone (E), DarE-DarA (EA), DarE-DarA-SAM (EAS), and DarE-SAM (ES) across the indicated labeling times. Data are shown as mean ± sd.

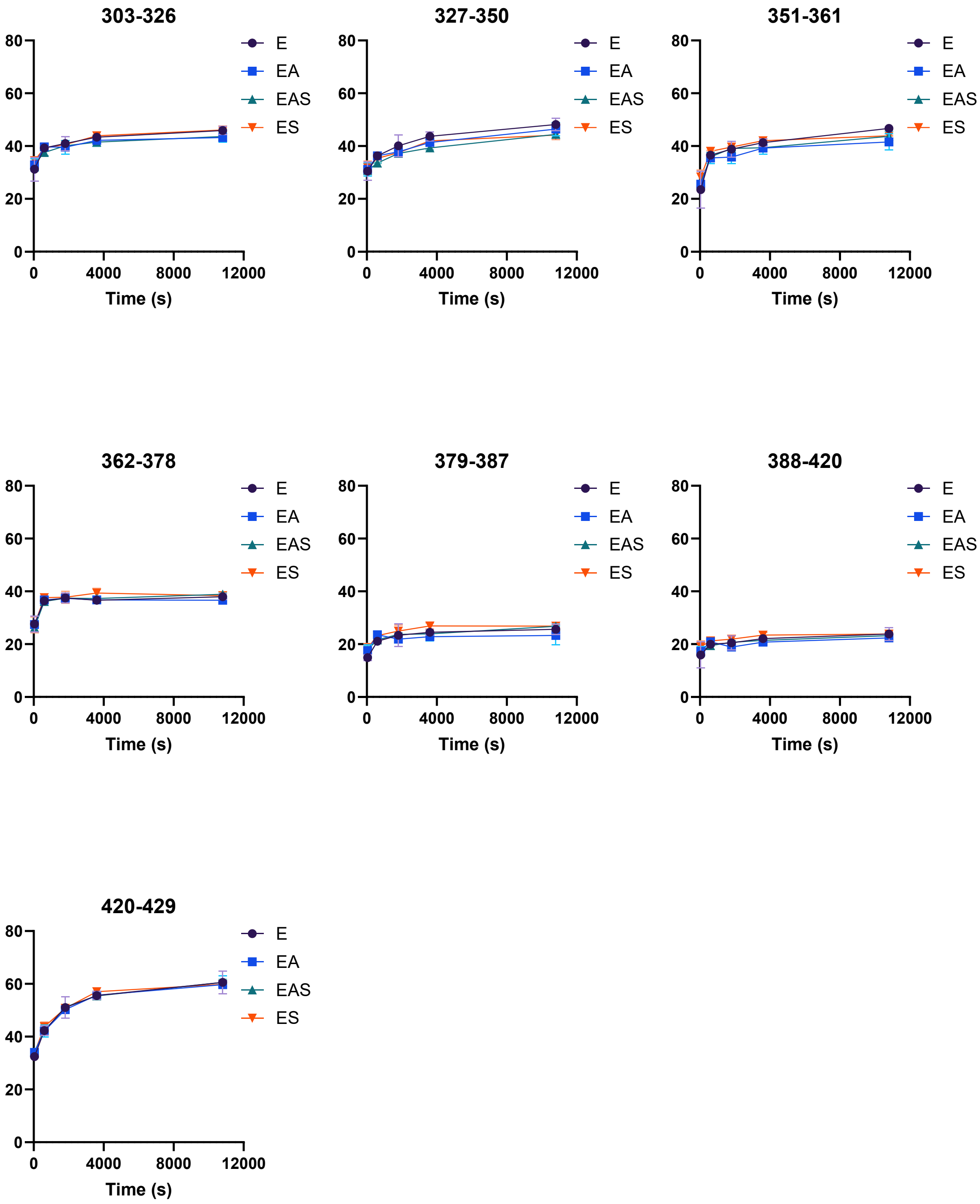


**Figure S4 (continued). Percentage deuterium uptake plots of DarE-derived peptides.** Deuterium uptake kinetics are shown for DarE-derived peptides. Percent deuterium uptake was measured for DarE alone (E), DarE-DarA (EA), DarE-DarA-SAM (EAS), and DarE-SAM (ES) across the indicated labeling times. Data are shown as mean ± sd.

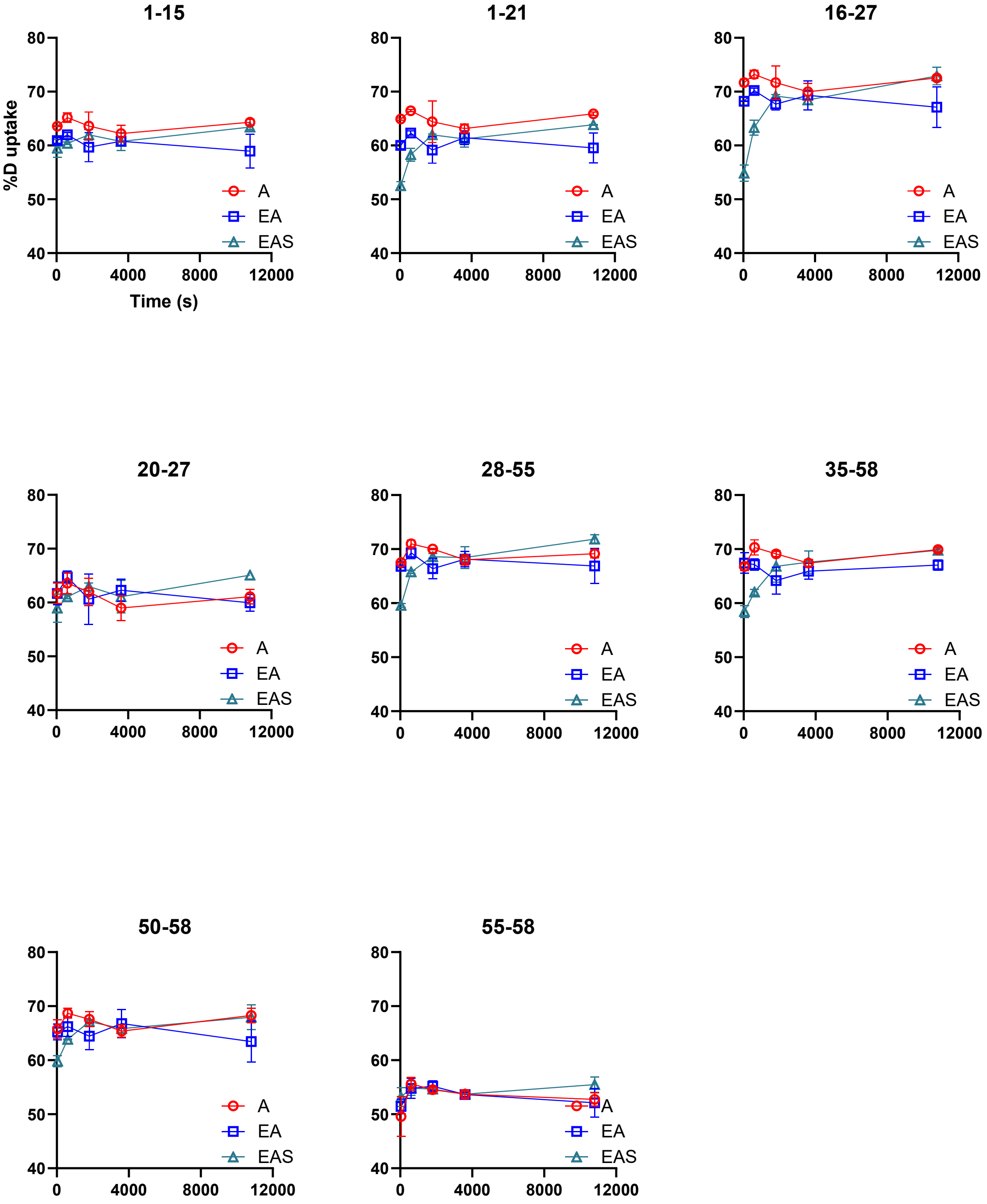


**Figure S5. Percentage deuterium uptake plot of DarA-derived peptides.** Deuterium uptake kinetics are shown for DarA-derived peptides. Percent deuterium uptake was measured for DarA alone (A), DarE-DarA (EA), and DarE-DarA-SAM (EAS) across the indicated labeling times. Data are shown as mean ± sd.

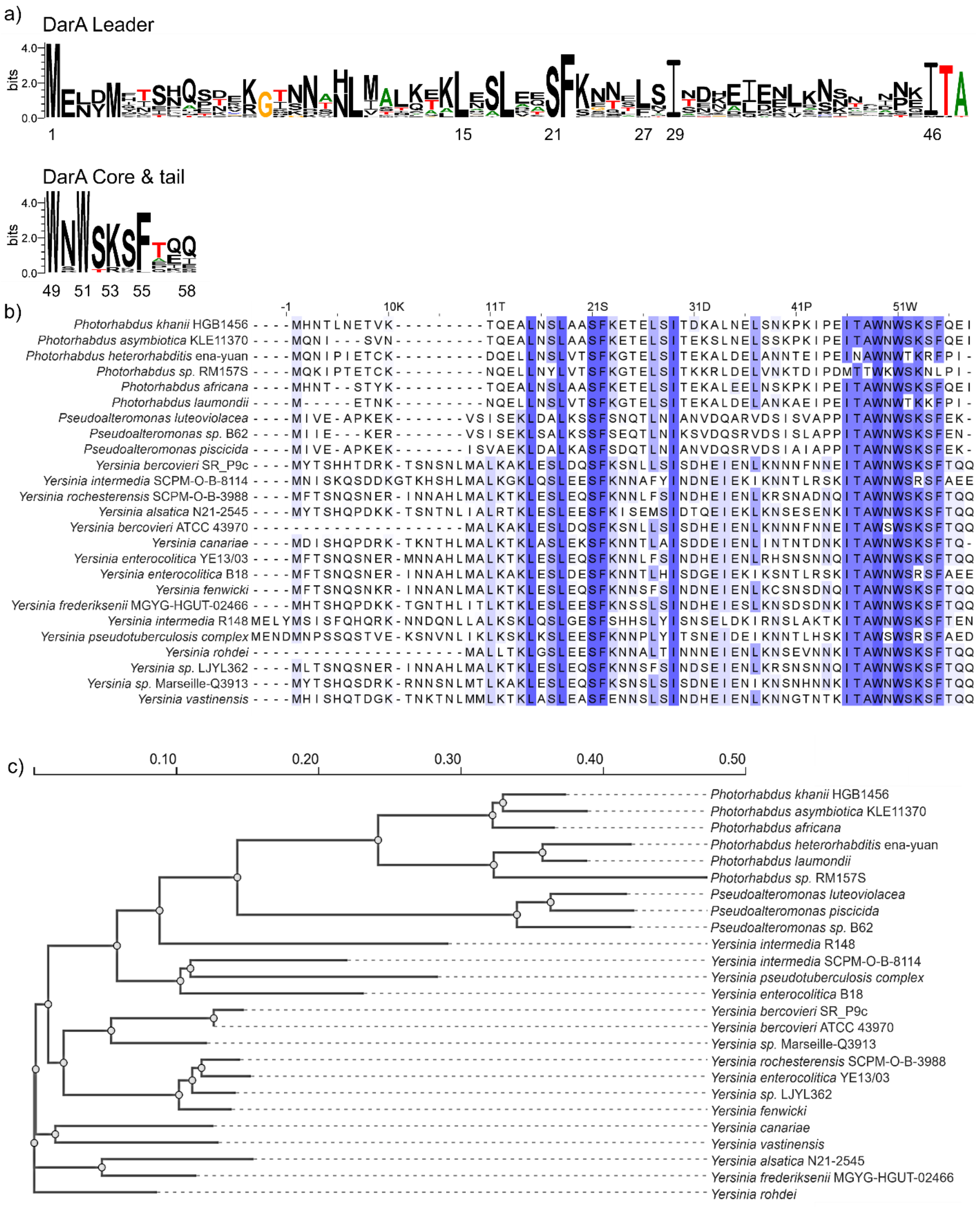


**Figure S6. Sequence conservation analysis of DarA homologs.** a) Sequence logo generated from aligned DarA homologs. b) Multiple sequence alignment of representative DarA homologs. Conserved residues are highlighted in blue. c) Phylogenetic tree constructed from the DarA sequences analyzed in this work.

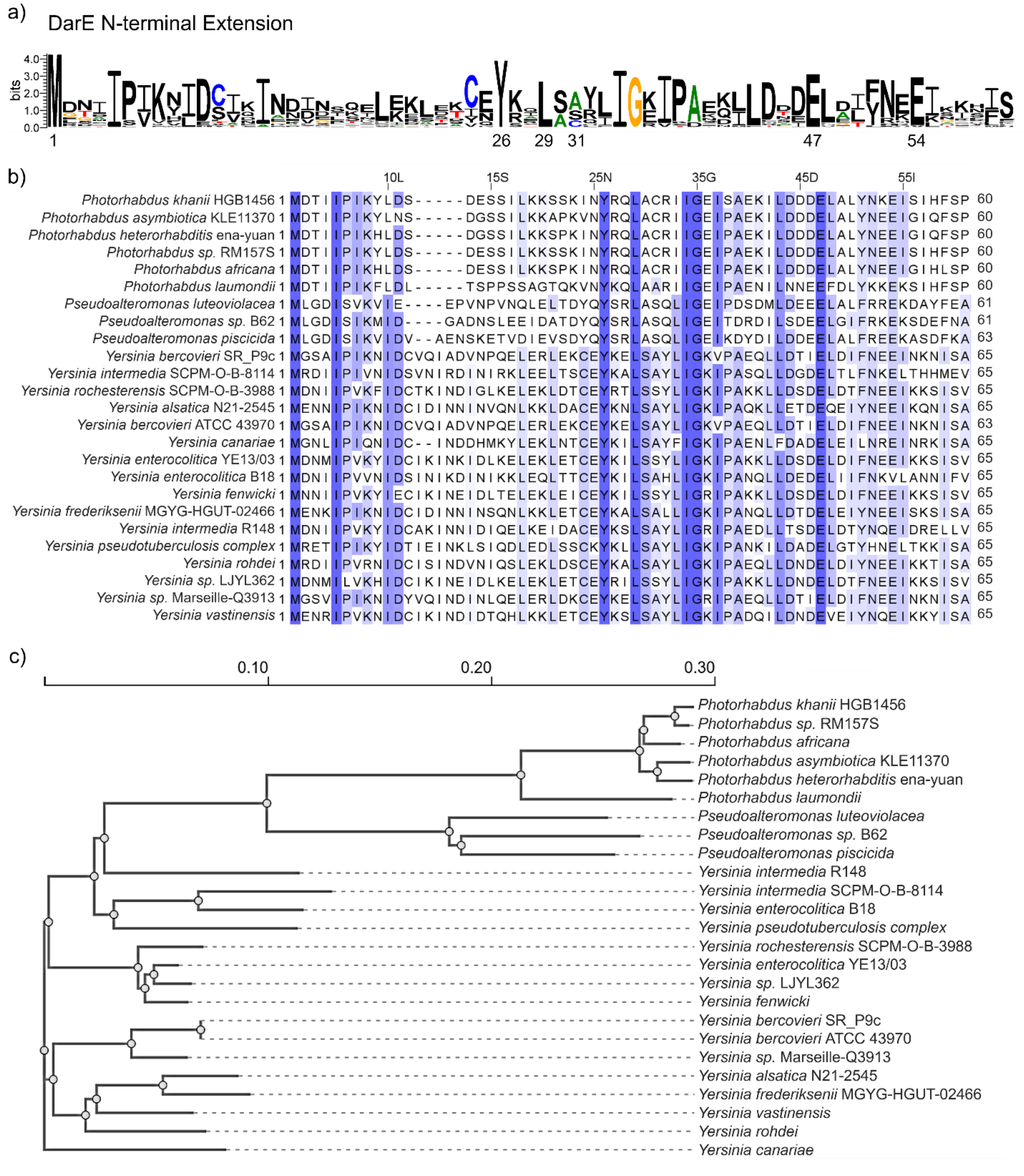


**Figure S7. Sequence conservation analysis of DarE homologs.** a) Sequence logo generated from the aligned N-terminal extension region across various DarE homologs. b) Multiple sequence alignment of representative DarE homologs. Conserved residues are highlighted in blue. c) Phylogenetic tree constructed from the DarE sequences analyzed in this work.

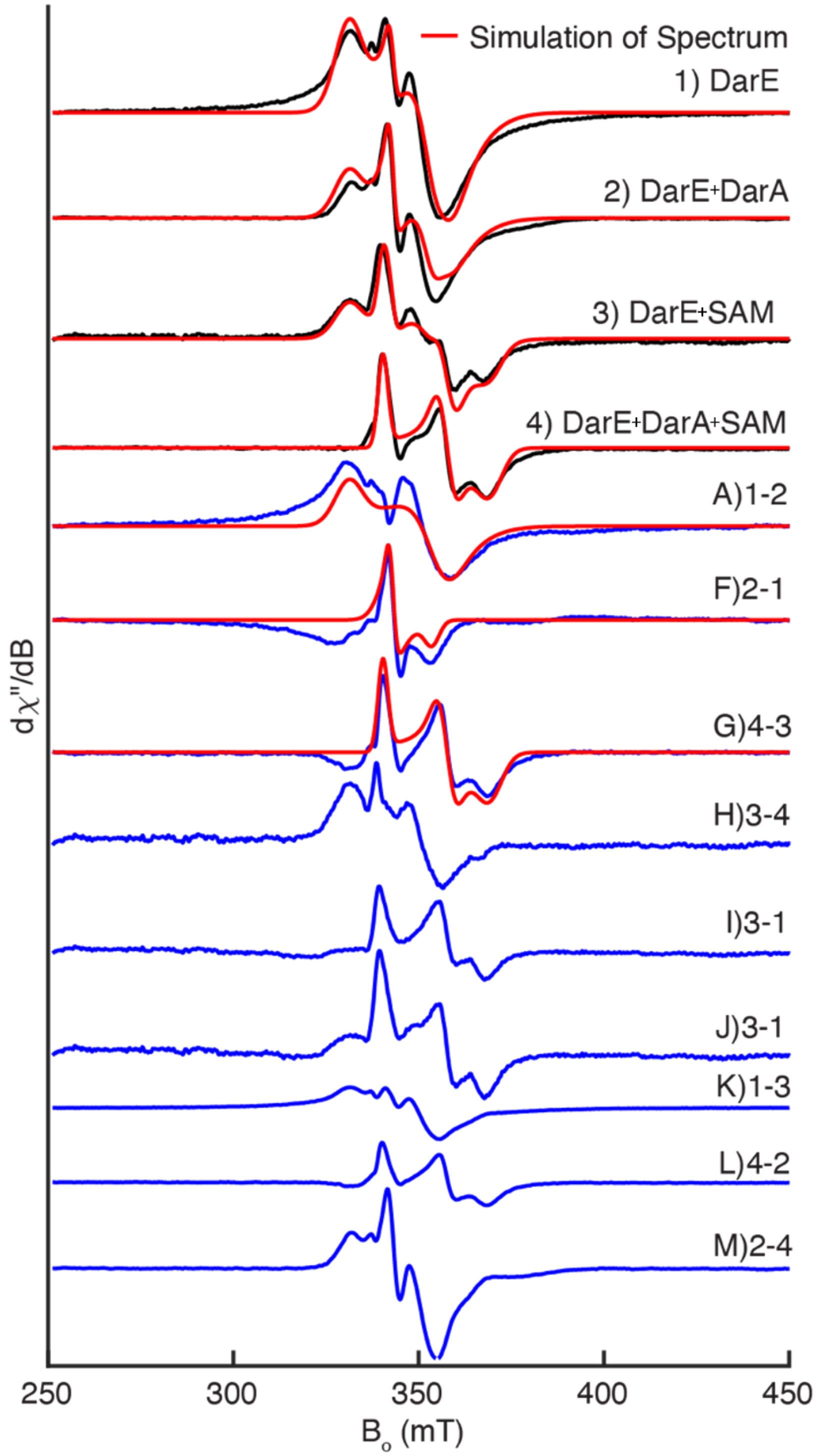


**Figure S8. CW EPR spectra and simulation of DarE complexes under sodium dithionite-reduced conditions.** Spectra correspond to 1) DarE alone, 2) DarE+DarA, 3) DarE+SAM, and 4) DarE+DarA+SAM complex, respectively. A)-J) are the pairwise difference spectra as indicated on sample sets. The lower traces (K-M) show individual simulated components used to reproduce the composite spectra, with the corresponding component assignments indicated at right. Experimental spectra are shown in black, the simulated data are shown in red, and subtracted spectra are shown in blue.

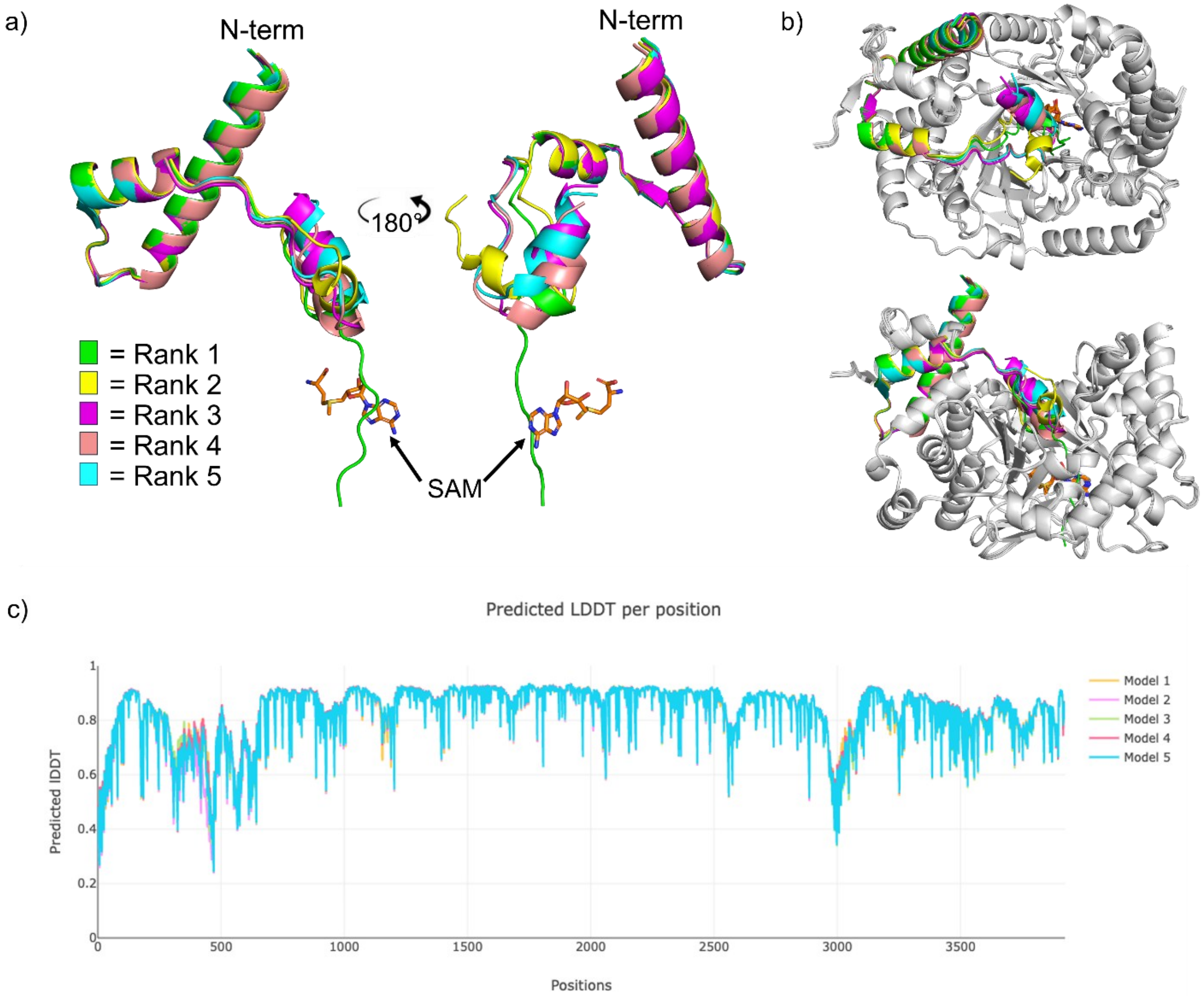


**Figure S9. Top 5 ranked OpenFold3 predicted EAS complex models.** a) Superposition of DarA from the five highest-ranked models reveals that the major structural variation occurs in the positioning of the DarA C-terminal core region. SAM is shown as orange sticks. b) The five predicted DarA conformations are shown in different colors and superimposed on DarE, shown in gray. c) Predicted lDDT (pLDDT) scores plotted by residue position for the five models.

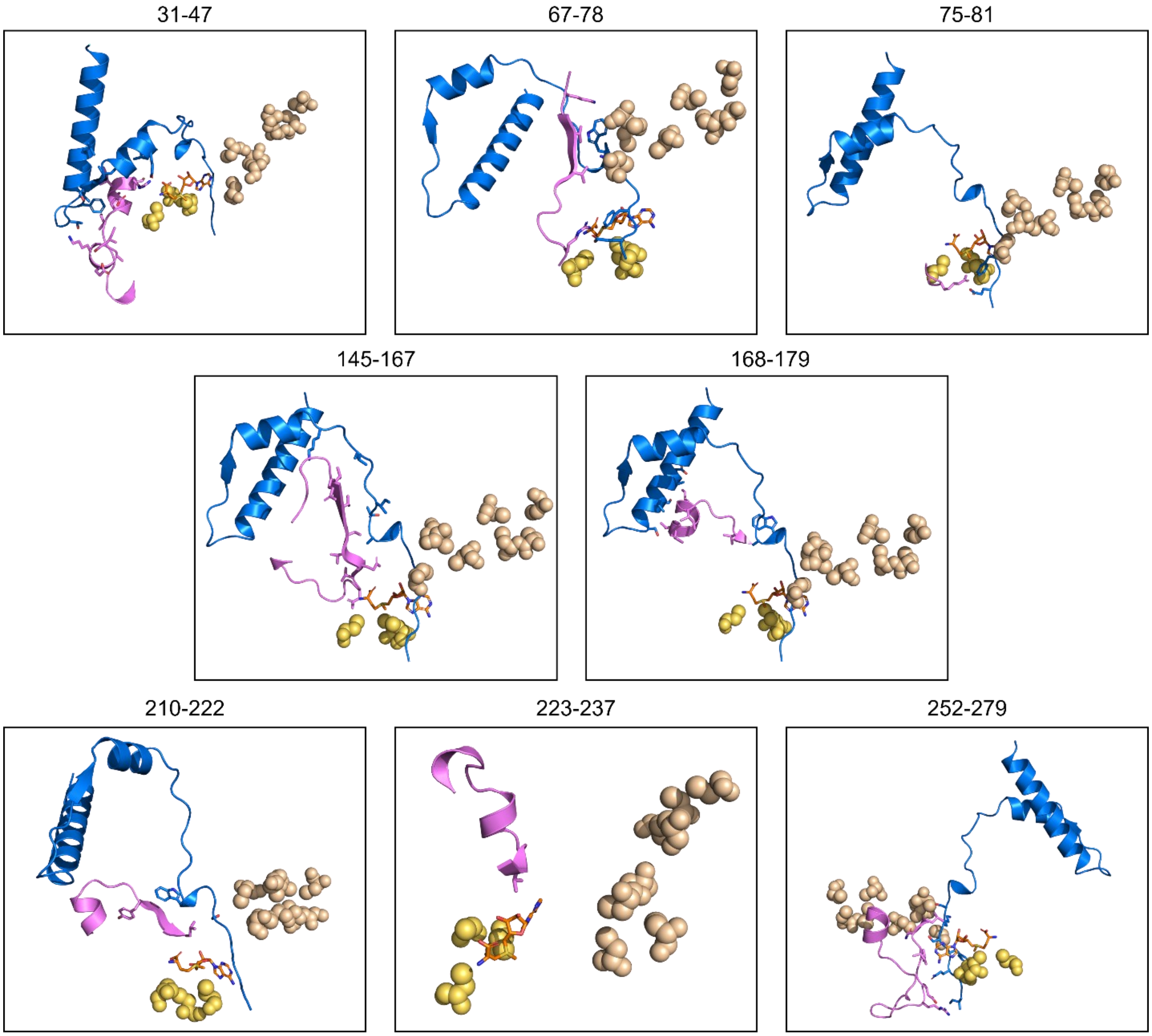


**Figure S10. Mapping HDX-MS-identified interactions onto AlphaMultimer-predicted complexes.** DarE-derived peptides that exhibited decreased deuterium uptake upon binding DarA and/or SAM are highlighted in pink on the predicted structural models. DarA is shown in blue, and SAM is shown as orange sticks. Cysteine residues coordinating the radical SAM [4Fe-4S] cluster are shown as yellow spheres, whereas cysteine ligands coordinating the AuxI and AuxII [4Fe-4S] clusters are shown as wheat-colored spheres.

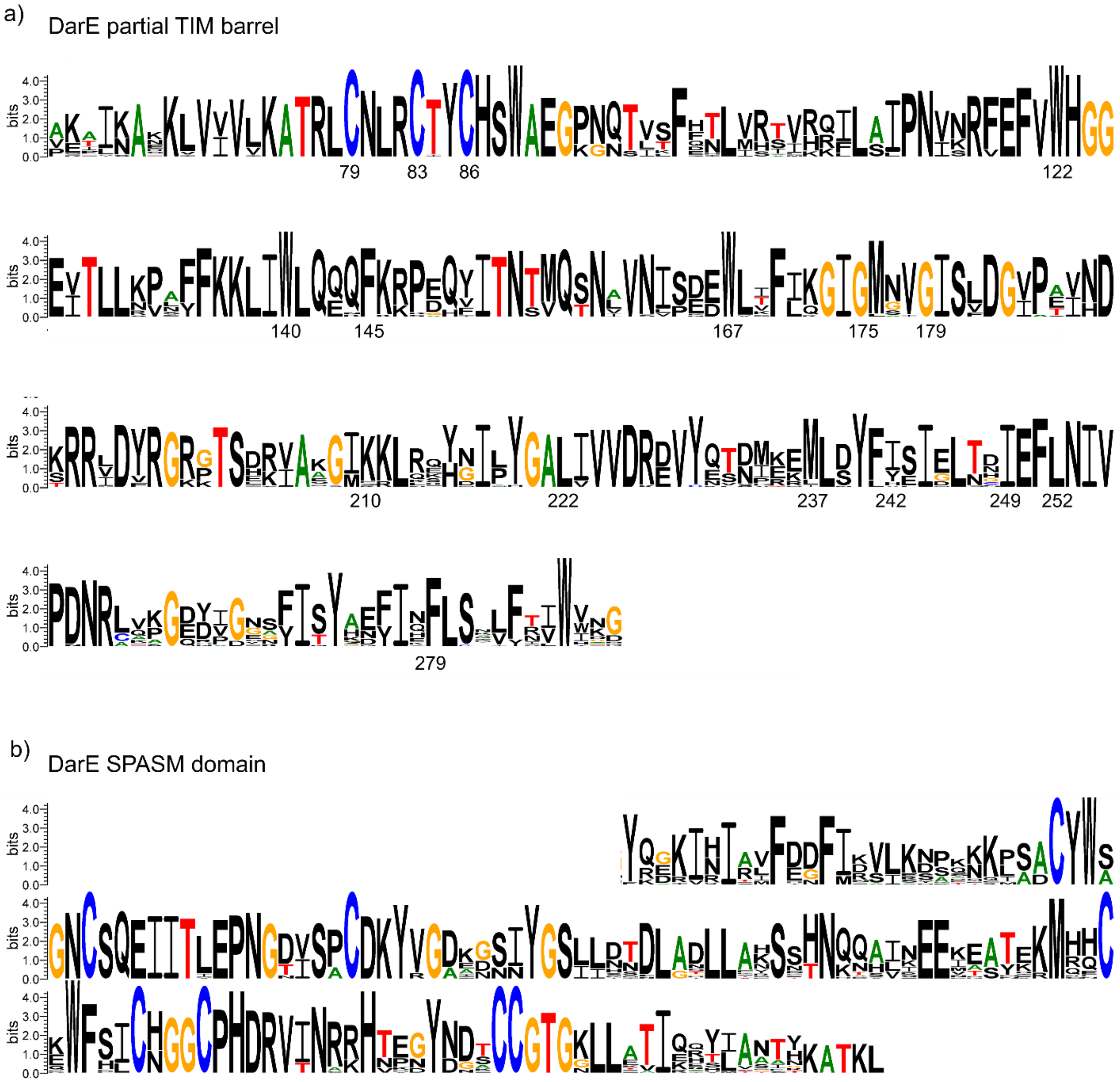


**Figure S11. Sequence conservation analysis of DarE homologs.** a) Sequence logo generated from the aligned partial TIM barrel across various DarE homologs. b) Sequence logo generated from the aligned SPASM domain across various DarE homologs.

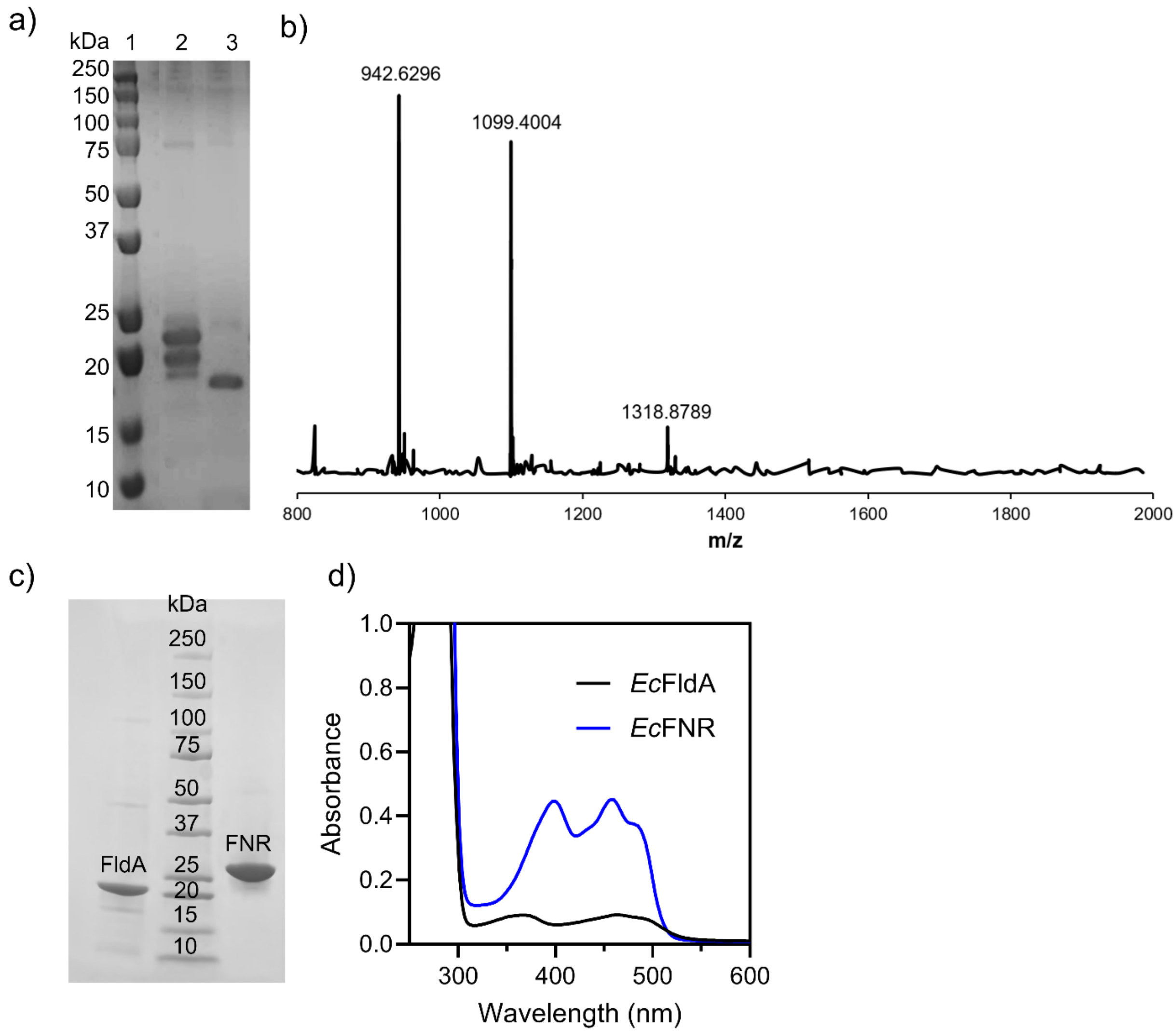


**Figure S12. Preparation and characterization of DarA and biological reductants.** a) SDS-PAGE of as-purified SUMO-DarA protein and ULP1 protease cleavage evaluation. Lane 1: Protein marker; Lane 2: as-purified SUMO-DarA protein; Lane 3: as-purified SUMO-DarA protein+ULP1 protease. b) LC-MS confirmed the production of DarA. c) SDS-PAGE of as-purified *E. coli* flavodoxin (*Ec*FldA) and *E. coli* flavodoxin $NADP^+$ reductase (*Ec*FNR). d) UV-vis spectra of the cofactor-loaded *Ec*FldA and *Ec*FNR
